# Amphiphilic proteins coassemble into multiphasic condensates and act as biomolecular surfactants

**DOI:** 10.1101/2021.05.28.446223

**Authors:** Fleurie M. Kelley, Bruna Favetta, Roshan M. Regy, Jeetain Mittal, Benjamin S. Schuster

## Abstract

Cells contain membraneless compartments that assemble due to liquid-liquid phase separation, including biomolecular condensates with complex morphologies. For instance, certain condensates are surrounded by a film of distinct composition, such as Ape1 condensates coated by a layer of Atg19, required for selective autophagy in yeast. Other condensates are multiphasic, with nested liquid phases of distinct compositions and functions, such as in the case of ribosome biogenesis in the nucleolus. The size and structure of such condensates must be regulated for proper biological function. We leveraged a bio-inspired approach to discover how amphiphilic, surfactant-like proteins may contribute to the structure and size regulation of biomolecular condensates. We designed and examined families of amphiphilic proteins comprising one phase-separating domain and one non-phase separating domain. In particular, these proteins contain the soluble structured domain glutathione S-transferase (GST) or maltose binding protein (MBP), fused to the intrinsically disordered RGG domain from P granule protein LAF-1. When one amphiphilic protein is mixed in vitro with RGG-RGG, the proteins assemble into enveloped condensates, with RGG-RGG at the core, and the amphiphilic protein forming the surface film layer. Importantly, we found that MBP-based amphiphiles are surfactants and control droplet size, with increasing surfactant concentration resulting in smaller droplet radii. In contrast, GST-based amphiphiles at increased concentrations co-assemble with RGG-RGG into multiphasic structures. We propose a mechanism for these experimental observations, supported by molecular simulations of a minimalist model. We speculate that surfactant proteins may play a significant role in regulating the structure and function of biomolecular condensates.

## Introduction

The intracellular environment is like a complex emulsion. This paradigm originated more than a century ago but is enjoying a renaissance, with recent discoveries revealing the important role of liquid-liquid phase separation (LLPS) in biology^1–3^. LLPS of proteins and nucleic acids underlies the formation of membraneless organelles, alternatively called biomolecular condensates, which are distinct intracellular compartments that lack a delimiting membrane^2,3^. Biomolecular condensates contribute to numerous cell functions, including stress response, gene regulation, and signaling^4^. Conversely, aberrant phase separation due to mutations and age-related processes is implicated in diseases such as neurodegeneration and cancer^5^. Deciphering the rules of self-assembly of biomolecular condensates has therefore emerged as a promising avenue for elucidating fundamental principles of biological structure, function, and dysfunction.

Despite significant recent progress in understanding the biophysics of biomolecular condensates, many open questions remain^6^. One key question is what molecular phenomena govern the spontaneous assembly of condensates with core-shell or multiphasic structures. Another important question is how cells tune the size of biomolecular condensates. Here, we sought to gain insight into both questions by examining how amphiphilic, surfactant-like proteins contribute to the self-assembly and regulation of biomolecular condensates. Amphiphiles are typically defined as molecules comprising separate hydrophilic and hydrophobic parts. Here, we note the etymology (in Greek, “amphi” means both and “philia” means friendship or love) and use the term amphiphile to describe proteins comprising one domain that has affinity for biomolecular condensates and one domain that has affinity for the dilute phase.

Surfactants are substances, generally amphiphiles, that adsorb to interfaces and decrease interfacial tension. Extracellularly, pulmonary surfactant lining the alveoli plays a vital role in lung physiology by reducing the work of breathing^7,8^. However, the role of surfactants in the emulsion-like intracellular milieu is just beginning to be explored^9^. In biological systems, the surfactant-like protein Ki-67 prevents individual chromosomes from coalescing during early stages of mitosis by forming a repulsive molecular brush layer^10^. Some biomolecular condensates are reminiscent of surfactant-laden emulsions, although their physical chemistry remains to be elucidated. For instance, Atg19 forms a thin surface layer surrounding Ape1 condensates that is necessary for selective autophagy of Ape1 in yeast^11^. Inspired by such examples, we hypothesized that a minimal system comprising surfactant-like proteins interacting with phase-separating proteins could recapitulate enveloped condensate structures observed in nature.

Moreover, condensates exhibit a variety of multiphase and multilayer structures underpinning their biological functions. Bre1 assembles as a shell surrounding Lge1 condensates, generating a catalytic condensate that functions to accelerate ubiquitination of histone H2B in yeast^12^. The nucleolus is comprised of coexisting liquid phases of differing interfacial tensions^13^, while P granules contain coexisting liquid and gel phases^14^. Stress granules^15^, nuclear speckles^16^, paraspeckles^17^, and reconstituted polypeptide/RNA complex coacervates also exhibit core-shell structures sensitive to stoichiometry and competitive binding^18–21^. Functionally related condensates can remain in contact without coalescing, as in the case of stress granules and P-bodies^22^, or P granules and Z granules^23^. We asked whether amphiphilic proteins could contribute to the complex morphologies of biomolecular condensates that have been observed within cells, just as synthetic amphiphiles and surfactant systems exhibit rich structures and phase behaviors^24^.

Surfactant-like proteins could have additional important functional consequences, including but not limited to modulating biomolecular condensate size, which in turn influences biochemical processes through condensate size-dependent effects on molecular concentrations and diffusion^25,26^. Biomolecular condensates are often observed in cells as multiple smaller droplets, rather than as a single larger droplet, even though the latter is expected to be thermodynamically favored. Recent studies have attributed the apparent metastability of biomolecular condensates in various contexts to surface charge^27^, cytoskeletal caging^28,29^, membrane association^30^, and exhaustion of available binding sites^31^; active processes can also maintain the emulsified, multi-droplet state in vivo^32^. An additional possibility, which we examine here, is that surfactant proteins may help stabilize biomolecular condensates.

To address these questions, we adopted a bottom-up, bioinspired approach, seeking to leverage a simplified system to shed light on the role of amphiphilic proteins in the self-assembly of biomolecular condensates. We designed amphiphilic proteins containing an intrinsically disordered region (IDR) fused to folded domains. The IDR is a phase-separating domain, whereas the folded domains are not. When one of these amphiphilic fusion proteins is mixed at low concentrations with the IDR alone, the two proteins assemble such that the amphiphilic protein forms a film that coats the IDR core. We demonstrate several extensions of this observation, including the important finding that condensate size can be controlled by varying surfactant protein concentration. Furthermore, when amphiphilic proteins with different folded domains are mixed together with the core IDR, we observe competition between different surfactant proteins for binding to the condensate interface. Interestingly, one family of amphiphilic proteins exhibits varied morphologies, including multiphasic condensates, and we map the rich concentration-dependent phase behavior. To gain mechanistic insight into these experimental observations, we present a minimalistic computational model that recapitulates the range of behaviors observed experimentally by varying the strength of interaction between domains. Our experiments and simulations suggest that amphiphile-condensate assembly is determined by the strength of interaction between the amphiphile and the IDR core, as well as interactions between the folded domain of the amphiphile. Taken together, this work illustrates the diverse interfacial phenomena that can arise from interactions between condensates and amphiphilic proteins, notably raising the possibility that surfactant proteins may play a significant role in regulating the structure and function of biomolecular condensates.

## Results

### Designing Amphiphilic Proteins to Act as Biological Surfactants

In prior work, we examined the partitioning of client proteins into biomolecular condensates^33^. We focused on the RGG domain from LAF-1, a prototypical arginine/glycine-rich intrinsically disordered protein (IDP) involved in P granule assembly in C. elegans^34–36^. Tandem repeats of the RGG domain (RGG-RGG) phase separate in vitro at concentrations >1 μM under physiological conditions. We found that RFP and GST-RFP are excluded from RGG-RGG condensates; RFP denotes red fluorescent protein and GST denotes glutathione S-transferase, which is widely used as a solubility-enhancing affinity tag in recombinant protein production^37^. In contrast, we observed that RFP fused to one or two RGG domains partitioned into RGG-RGG condensates. These results demonstrated that client partitioning into or exclusion from RGG-RGG condensates depends on the balance between the “RGG-philic” and “RGG-phobic” content of the client protein. Here, we hypothesized that a third, intermediate outcome is possible: that with the right balance, a client protein may localize to the interface between the condensate and dilute phases, coating the condensate and displaying surfactant-like behavior.

We therefore asked whether we could engineer amphiphilic proteins that assemble as a film on the surface of RGG-RGG condensates. We designed a family of amphiphilic fusion proteins containing a phase-separating domain (RGG-philic) fused to a non-phase separating domain (RGG-phobic) (SI Appendix Fig. 1). Two representative protein constructs from this family are MBP-GFP-RGG and GST-GFP-RGG (Fig. 1A). MBP denotes maltose binding protein, which like GST is a well-known affinity tag that enhances protein solubility^37–40^. GFP denotes enhanced green fluorescent protein with the monomerizing A206K mutation. By combining RGG, GST, MBP, and fluorescent protein domains, our aim was to generate amphiphilic fusion proteins with RGG-philic and RGG-phobic parts. We hypothesized that upon mixing the amphiphilic proteins with RGG-RGG, the poorly soluble RGG domain in the amphiphiles would interact with and orient towards the inner RGG-RGG condensate phase, while the RGG-phobic MBP and GST domains would interact with and orient towards the outer dilute phase (Fig. 1B).

**Figure 1:**
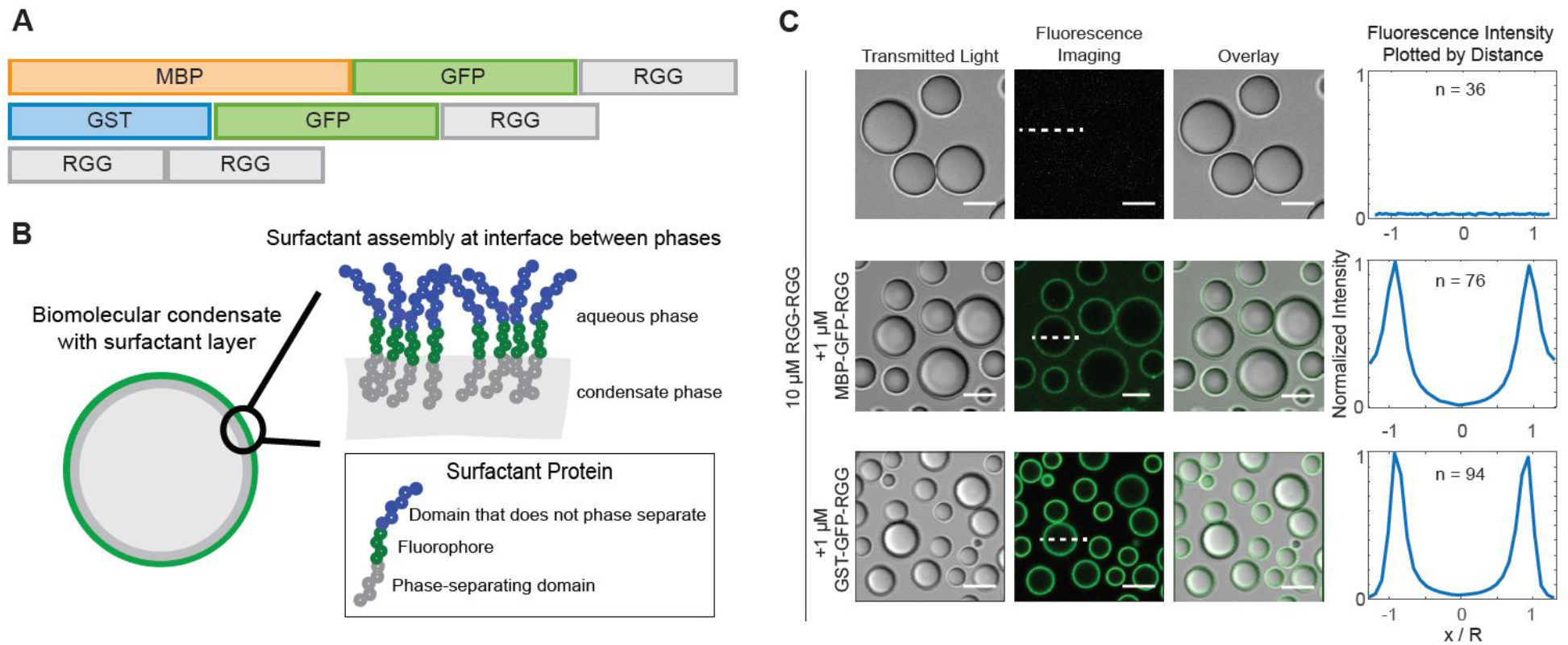
Amphiphilic proteins MBP-GFP-RGG and GST-GFP-RGG form a film around RGG-RGG. **A**, Schematic domain diagram for MBP-GFP-RGG, GST-GFP-RGG, and RGG-RGG. Length is proportional to number of base pairs. **B**, Schematic model of the interaction between amphiphilic proteins and RGG-RGG. The amphiphilic protein is depicted at the interface between the two phases, surrounding the RGG-RGG core, with the phase-separating RGG domain facing the RGG-RGG core and the non-phase-separating GST or MBP domain facing the aqueous phase. **C**, Transmitted light and fluorescence imaging of the film formed by mixing RGG-RGG and MBP-GFP-RGG or GST-GFP-RGG in a 10:1 concentration ratio (10 μM RGG-RGG, 1 μM MBP-GFP-RGG or GST-GFP-RGG). For this and subsequent figures, buffer was 150 mM NaCl, 20 mM Tris, pH 7.5. Right: Fluorescence intensities were quantified as line profiles across individual condensates, which were normalized and averaged. n, number of condensates. Scale bars, 5 μm.

### Amphiphilic Proteins Coat Biomolecular Condensates, Resembling Surfactants

To test our hypothesis, we mixed RGG-RGG with the amphiphilic proteins GST-GFP-RGG or MBP-GFP-RGG in physiological buffer and used microscopy to observe the resulting assemblies. In these binary mixtures, RGG-RGG phase separated into spherical protein droplets, as expected. Remarkably, MBP-GFP-RGG formed green rings around the RGG-RGG condensates, as observed by confocal fluorescence microscopy (Fig. 1C). Similarly, GST-GFP-RGG mixed with RGG-RGG formed green rings around the RGG-RGG condensates. We analyzed fluorescence intensity line profiles drawn radially across condensates in the confocal images, showing localization of GFP mainly at the condensate surface, as opposed to within or outside the condensates (Fig. 1C). Quantification of the partitioning of both surfactant-like proteins indicates a subtle difference. While GST-GFP-RGG localized almost entirely to the interface, MBP-GFP-RGG exhibits an elevated concentration in the dilute phase (ratio of concentrations at interface: condensate: buffer for GST-GFP-RGG is 1: 0.05: 0.01 and for MBP-GFP-RGG is 1: 0.05: 0.3). This suggests a potentially higher affinity between the GST-GFP-RGG protein and RGG-RGG condensate when compared to MBP-GFP-RGG. In both cases, these data demonstrate that the amphiphilic proteins adsorb to and form a layer at the condensate surface, reminiscent of surfactant-stabilized emulsions. This surfactant-like assembly was observed even upon shuffling the sequence of the RGG domains used to form the droplet core, suggesting that the adsorption does not depend on the precise sequence of the phase-separating IDP (SI Appendix Fig. 2).

To confirm the mechanism by which the amphiphilic proteins adsorb to the interface, we examined how the fluorescence localization changes when the RGG-philic and RGG-phobic regions of the amphiphilic protein are uncoupled. We therefore included a tobacco etch virus (TEV) protease recognition site immediately N-terminal to the RGG domain of MBP-GFP-RGG, so that upon treatment with TEV protease, MBP-GFP would be liberated from RGG (Fig. 2A). We hypothesized that when TEV protease is added to RGG-RGG condensates enveloped by an MBP-GFP-RGG film, the cleaved RGG domain from the surfactant protein would partition into the RGG-RGG condensates, whereas the MBP-GFP would partition into the surrounding dilute phase, so the fluorescence intensity of the ring would decrease (Fig. 2B). Indeed, using time-lapse confocal microscopy, we observed the intensity of the fluorescent ring rapidly diminish following TEV protease addition (Fig. 2C-D). The fluorescence intensity of the rings, f(t), decays approximately exponentially with time. The data is consistent with first-order kinetics, *f*(*t*) = *a* * *e^−k*t^*, which is expected for an enzyme-catalyzed reaction in which the enzyme is not saturated by the substrate ([GST-GFP-RGG] of 1 μM ≪ Michaelis constant K_m_ of TEV ~ 61 μM^41^). As a control, we performed the same experiment without TEV protease to account for photobleaching and observed only slow, linear decrease in fluorescence intensity (SI Appendix Fig. 3). Collectively, these results support the model that MBP-GFP-RGG and related proteins localize to the condensate interface due to their amphiphilic nature.

**Figure 2:**
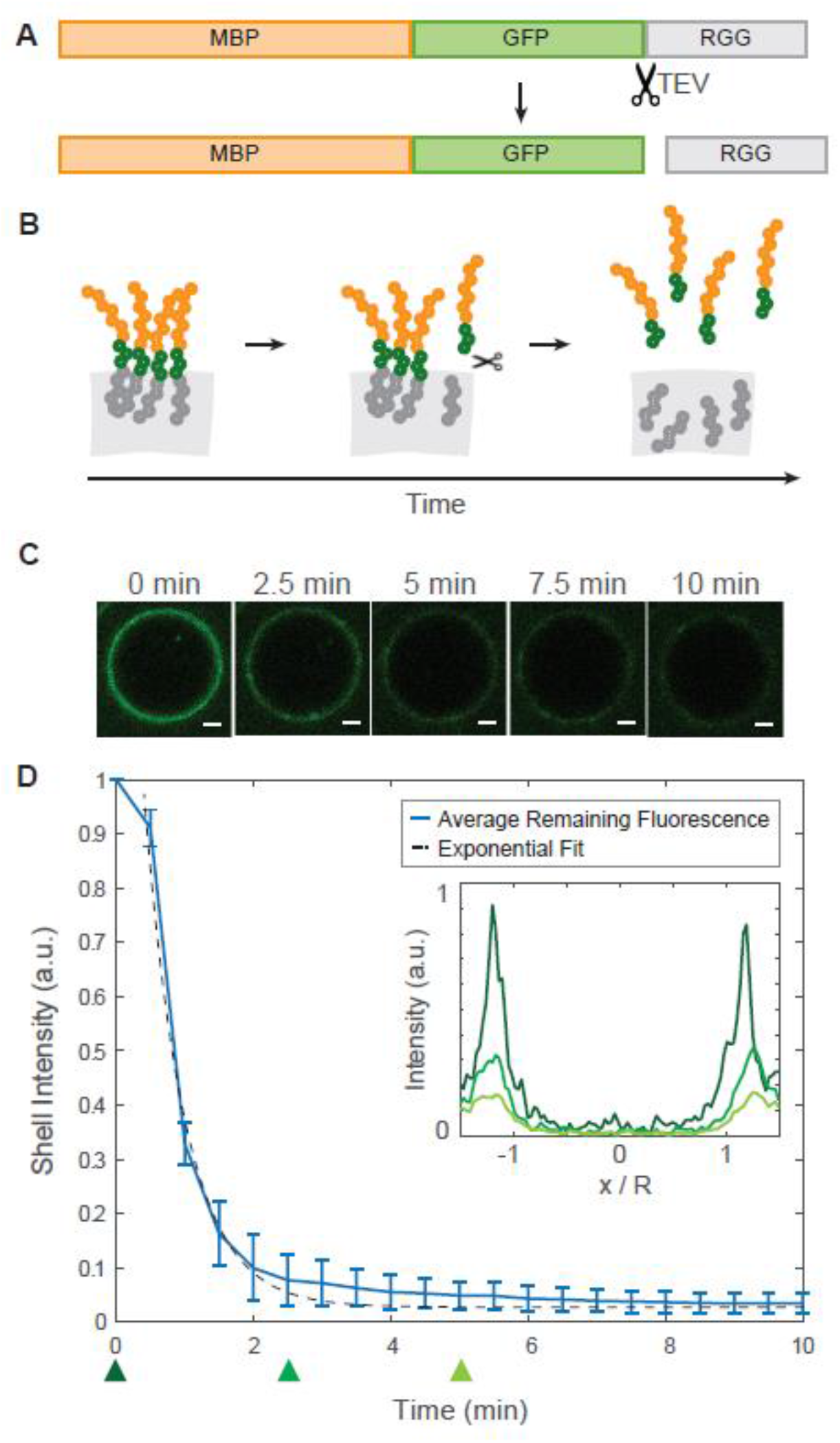
Desorption of surface-associated amphiphilic protein triggered by proteolytically cleaving the dissimilar domains. **A**, Schematic depicting the location of the TEV protease cleavage site inserted into MBP-GFP-RGG, between the GFP and RGG domains. **B**, Schematic model of the result of adding TEV protease to the RGG-RGG and MBP-GFP-RGG system. Cleavage of the amphiphilic protein results in the release of the fluorescently tagged MBP domain into the aqueous solution and retention of the RGG domain by the condensed phase. **C**, Representative droplet images at 0, 2.5, 5, 7.5 and 10 minutes after TEV protease addition. Scale bars, 1 μm. **D**, Total fluorescence intensity of the shell region decreases over time with the release of MBP-GFP following cleavage of MBP-GFP-RGG by TEV protease, which was added at t = 0 (plot shows average of n = 87 droplets). Inset, normalized average fluorescence intensity line profiles at time 0 (dark green), 2.5 minutes (medium green) and 5 minutes (light green). 5 mM DTT was added to the buffer in accordance with recommended TEV protease protocols.

### Surfactant-like Proteins Control Condensate Size

In conducting the aforementioned TEV protease experiments, we observed that removing the protein film from around the RGG-RGG condensates triggered droplet fusion events. We sought to further understand this result, asking whether the amphiphilic proteins can control condensate size by acting as biological surfactants and stabilizing condensates against fusion. We therefore measured droplet sizes upon mixing RGG-RGG with various concentrations of the amphiphilic protein MBP-GFP-RGG (Fig. 3A). As the concentration of MBP-GFP-RGG increased, the droplet sizes decreased significantly, as observed by brightfield and confocal microscopy (Fig. 3B). This would be expected for surfactants in a surfactant-poor regime, where there is no excess of emulsifier^42^. Since the condensates displayed a wide range of sizes for any given MBP-GFP-RGG surfactant concentration, we plotted histograms of droplet size, further revealing a shift from larger condensates in the absence of surfactant to numerous smaller condensates as surfactant concentration increases (Fig. 3C). Interestingly, size distribution appears bimodal in several conditions, which is reminiscent of observations in nanocrystal growth where the smaller-size mode corresponds to primary nucleation, and the larger-size mode to the nanoparticles growing by aggregation^43^. Total condensate volume is approximately conserved across samples with differing surfactant concentrations (+/- 19%), supporting the theory that surfactant proteins do not inhibit phase separation but rather stabilize droplet size. Motivated by studies on synthetic triblock polymer surfactants, comprised of a hydrophobic block sandwiched between hydrophilic blocks, we wondered how our results may differ if the RGG domain were placed as the middle block of a surfactant protein. We therefore switched the order of the fluorescent and phase-separating domains, forming MBP-RGG-GFP. We confirmed that MBP-RGG-GFP also coats RGG-RGG condensates, appearing as fluorescent rings under confocal microscopy. Similar to MBP-GFP-RGG, increasing concentrations of MBP-RGG-GFP resulted in decreased RGG-RGG droplet size (Fig. 3D-F).

**Figure 3:**
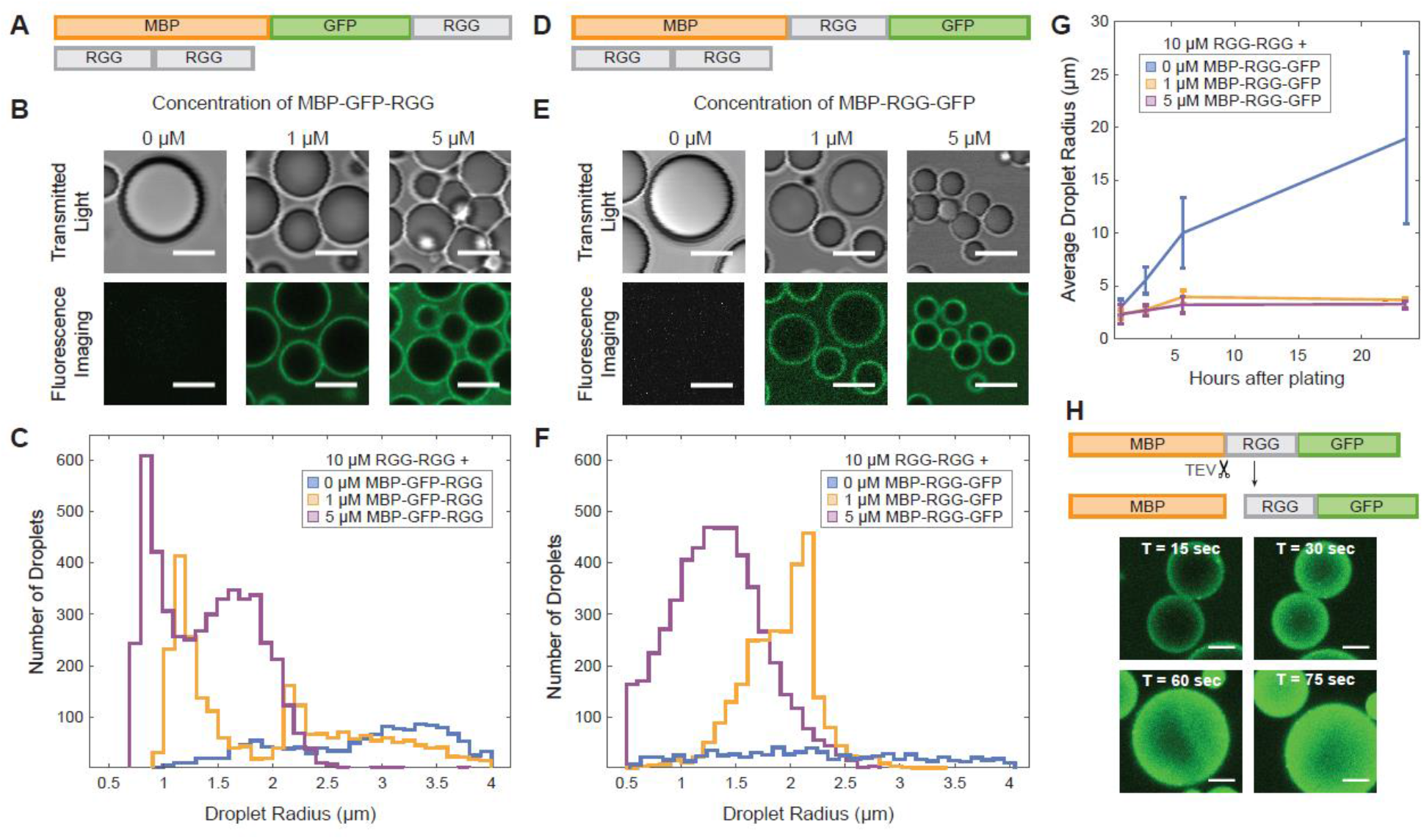
Amphiphilic proteins MBP-GFP-RGG and MBP-RGG-GFP act as surfactants by stabilizing small condensates. **A**, Domain schematics depicting surfactant protein MBP-GFP-RGG and core protein RGG-RGG. **B**, Transmitted light and confocal fluorescence imaging of the film formed by mixing RGG-RGG (10 μM) with different concentrations of MBP-GFP-RGG. Droplet size becomes smaller as MBP-GFP-RGG concentration increases. Scale bars, 5 μm. **C**, Histogram depicting significant shift from a small number of large condensates when no surfactant protein is added, to a large number of small condensates with increasing surfactant protein concentration (p < 0.005 based on one-way ANOVA followed by Tukey’s post hoc test). **D**, Schematic depicting MBP-RGG-GFP and RGG-RGG. **E**, Micrographs depict progressively smaller droplet sizes with increasing concentrations of MBP-RGG-GFP, similar to the result for MBP-GFP-RGG. Scale bars, 5 μm. **F**, Histogram depicting shift in number and size of condensates with increasing concentrations of MBP-RGG-GFP (p < 0.005 based on one-way ANOVA followed by Tukey’s post hoc test). **G**, Surfactant stabilization of droplet size is sustained after 3, 6, and 24 hours (p < 0.05 based on one-way ANOVA followed by Tukey’s post hoc test). **H**, TEV protease cleaves the MBP-RGG-GFP construct between MBP and RGG, releasing MBP and causing partitioning of RGG-GFP into the RGG-RGG condensates. Rapid fusion events can be observed upon disassembly of the MBP-RGG-GFP layer. Scale bars, 1 μm.

Using this system, we also measured how RGG-RGG droplet size changes over time with or without MBP-RGG-GFP surfactant. Without a stabilizing mechanism, and provided that droplets do not age, condensates would be expected to grow over time due to coalescence and Ostwald ripening^44^. We examined droplet size at timepoints of 1, 3, 6, and 24 hours, finding that without surfactant, average droplet size indeed grew significantly over time (from average radius of 3.0 μm at hour 1 to 19.0 μm at hour 24). In stark contrast, we observed only a slight size increase for the surfactant-coated condensates (from average radius of 2.3 μm at hour 1, to 3.7 and 3.3 μm at hour 24, for 1 μM and 5 μM MBP-RGG-GFP, respectively). These data illustrate the lasting stabilizing effect of the surfactant layer (Fig. 3G). To explore the implications of these results, we further examined how droplet size changes upon triggered removal of the surfactant layer. We added TEV protease to RGG-RGG condensates coated with MBP-x-RGG-GFP, where x denotes a TEV protease recognition sequence, and we measured the change in cross-sectional area of the condensates over time. When the TEV protease cut site is placed such that MBP is liberated alone and GFP remains attached to RGG, TEV protease treatment causes the green fluorescence to relocalize from the interface to the interior of the condensates (Fig. 3H). Strikingly, removal of the surfactant layer triggered rapid fusion events, leading to a significant increase in droplet size. A majority of fusion events and size increase occurred within the first five minutes after TEV addition (SI Appendix Fig. 4), after which droplet size stabilized. As a control, we added TEV to a sample containing only RGG-RGG. Though scattered fusion events occurred due to the creation of flow, the average droplet size did not change significantly. Overall, our results suggest that condensate size can be controlled by surfactant proteins, dependent on surfactant concentration and tunable via stimuli that manipulate the interfacial properties.

### Rich Phase Behavior with Increasing Concentrations of GST-GFP-RGG

Next, we investigated the impact of altering the stoichiometry of the system. The experiments in which we varied the concentrations of MBP-GFP-RGG and MBP-RGG-GFP suggested that the amphiphilic protein can accumulate in the continuous phase as the surface becomes saturated (Fig. 3B,E). An alternative scenario, however, is that at high concentrations, the amphiphilic protein could itself phase separate, resulting in multiphasic behavior. A growing body of literature supports the existence and important role of multiphasic behavior in biological^13,14,45^ and polymer-based systems^46^. We therefore considered whether this occurs in our system, focusing on GST-GFP-RGG based on its lower partition coefficient in the continuous phase as compared to the MBP-based surfactants (Fig. 1C). For these experiments we used buffer consisting of 150 mM NaCl, 20 mM Tris, pH 7.5 with no reducing agent added.

Remarkably, we observed rich phase behavior in a system of varying concentrations of RGG-RGG plus GST-GFP-RGG. The various assemblies can be categorized into three regimes. As described previously, at low concentration of GST-GFP-RGG relative to RGG-RGG, we observed that RGG-RGG condensates are coated with a film of GST-GFP-RGG (Fig. 4A, left). With increasing relative concentration of GST-GFP-RGG to RGG-RGG, a transition occurs to a second regime where GST-GFP-RGG phase separates and forms puncta that partially engulf the core droplet (Fig. 4A, middle). Upon still further increasing the relative concentration of GST-GFP-RGG to RGG-RGG, the two proteins assemble into a multiphasic system where a continuous GST-GFP-RGG phase envelops RGG-RGG droplets (Fig. 4A, right). The formation of a GST-GFP-RGG phase at high concentrations, whereas MBP-based proteins did not display such behavior, implies a higher strength of interactions between GST domains or between GST and the RGG-RGG core.

**Figure 4:**
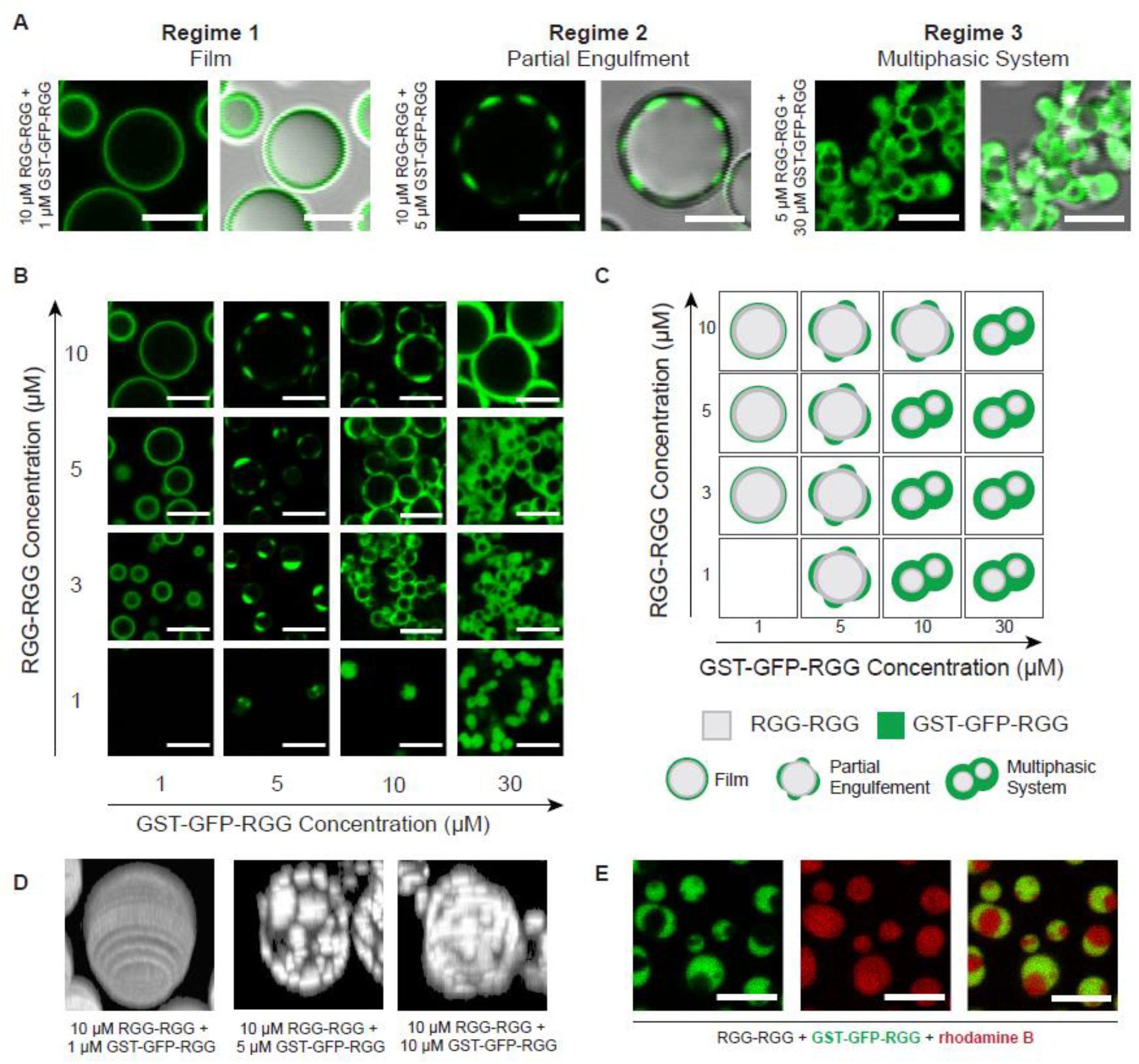
Rich phase behavior of the system depends on stoichiometry of RGG-RGG: GST-GFP-RGG. **A**, Classification system for the morphologies formed by mixing different ratios of RGG-RGG to GST-GFP-RGG. **B**, Representative images of condensates formed across concentrations of RGG-RGG and GST-GFP-RGG tested, depicting a range of phase behaviors. **C**, Application of classification system to the assemblies observed in the phase diagram, including film, partial engulfment, and multiphasic complexes. **D**, Three-dimensional volume renderings of representative condensates depicting (left) film, (middle) low coverage partial engulfment, (right) high coverage partial engulfment. **E**, Dual-color imaging of multiphasic system (incorporating rhodamine B) to confirm presence of RGG-RGG in engulfed phase. Scale bars, 5 μm.

To further explore this range of behaviors, we mapped the phase space by varying the concentrations of both RGG-RGG and GST-GFP-RGG (Fig. 4B). When 1 μM GST-GFP-RGG was added to 3, 5, or 10 μM RGG-RGG, we observed films of GST-GFP-RGG coating the RGG-RGG condensates, agreeing with our prior observations. When the concentration of GST-GFP-RGG was increased to 5 μM, we observed the formation of partially engulfed RGG-RGG droplets. At a GST-GFP-RGG concentration of 10 μM, we observed partial engulfment at high RGG-RGG concentration (10 μM), shifting to multiphasic behavior at lower RGG-RGG concentration. Finally, increasing the concentration of GST-GFP-RGG further to 30 μM resulted in a greater volume of the GST-GFP-RGG phase. We can schematically summarize the phase diagram based on these qualitative categorizations for each concentration pair (Fig. 4C).

Next, we generated 3-dimensional (3D) reconstructions of condensates from confocal z-stacks to show the degree of coverage of GST-GFP-RGG surrounding the RGG-RGG phase. In the first regime, with low GST-GFP-RGG concentration, the 3D reconstruction showed a homogenous fluorescent film surrounding the RGG-RGG condensate (Fig. 4D, left panel). At higher concentrations of GST-GFP-RGG, the 3D reconstructions revealed small GST-GFP-RGG condensates decorating larger core RGG-RGG condensates (Fig. 4D, middle panel). Still further increase of GST-GFP-RGG concentration resulted in increased surface coverage by the GST-GFP-RGG phase (Fig. 4D, right panel). Next, to better visualize the multiphasic regime at relatively low RGG-RGG and high GST-GFP-RGG concentrations, we added a small amount of rhodamine B to the system (Fig. 4E). The rhodamine strongly partitioned into both the RGG-RGG and GST-GFP-RGG phases (partition coefficients of 21 and 17, respectively, defined as the ratio of rhodamine fluorescence in each condensate phase to that of the dilute phase). This confirms that the voids observed in the GST-GFP-RGG phase are composed of condensed RGG-RGG, rather than buffer. Finally, we tested the liquidity of the GST-GFP-RGG phase compared to the RGG-RGG phase using fluorescence recovery after photobleaching (FRAP). We observed a striking difference: GST-GFP-RGG exhibited slow and incomplete recovery (approximately 10% recovery after 1 minute), compared to the faster and almost complete RGG-RGG recovery (approximately 70% after 1 minute; SI Appendix Fig. 5)^33^. This FRAP data suggests that GST-GFP-RGG has a higher viscosity compared to RGG-RGG.

### GST-based surfactants outcompete MBP-based surfactants at binding to the condensate interface

At first glance, GST-based and MBP-based amphiphiles exhibit similar behavior when either of the amphiphilic proteins is mixed at low concentration (1 μM) with RGG-RGG (10 μM), forming core condensates enveloped by thin surfactant films. However, we previously noted subtle differences in partitioning of GST-GFP-RGG vs. MBP-GFP-RGG in this regime. Whereas GST-GFP-RGG was highly localized to the droplet surface, MBP-GFP-RGG was somewhat less so, with measurable MBP-GFP-RGG fluorescence in the continuous phase (Figs. 1C and 3B) – suggesting these two amphiphilic proteins have different affinities to the RGG-RGG condensate surface. This notion is further supported by the differing behaviors of these proteins at increased concentrations, wherein GST-GFP-RGG phase separates preferentially at the RGG-RGG droplet surface while excess MBP-GFP-RGG accumulates in the continuous phase. This raises the question of how GST-based and MBP-based surfactants would behave upon mixing RGG-RGG concurrently with both GST-based and MBP-based surfactants. Based on our prior observations, we hypothesized that the two amphiphiles may compete for binding to the interface.

To begin, we needed to examine the effects on biomolecular assembly of varying the fluorescent protein. Both the GST-based and MBP-based surfactants displayed consistent behavior when GFP was replaced with the monomeric red fluorescent protein mCherry (Fig. 5A). Irrespective of fluorophore, individually the amphiphilic proteins (1 μM) colocalized to the surface of RGG-RGG (10 μM) droplets. This allowed us to assess interactions between the amphiphilic proteins, focusing on the role of GST vs. MBP, regardless of the fluorophore.

**Figure 5:**
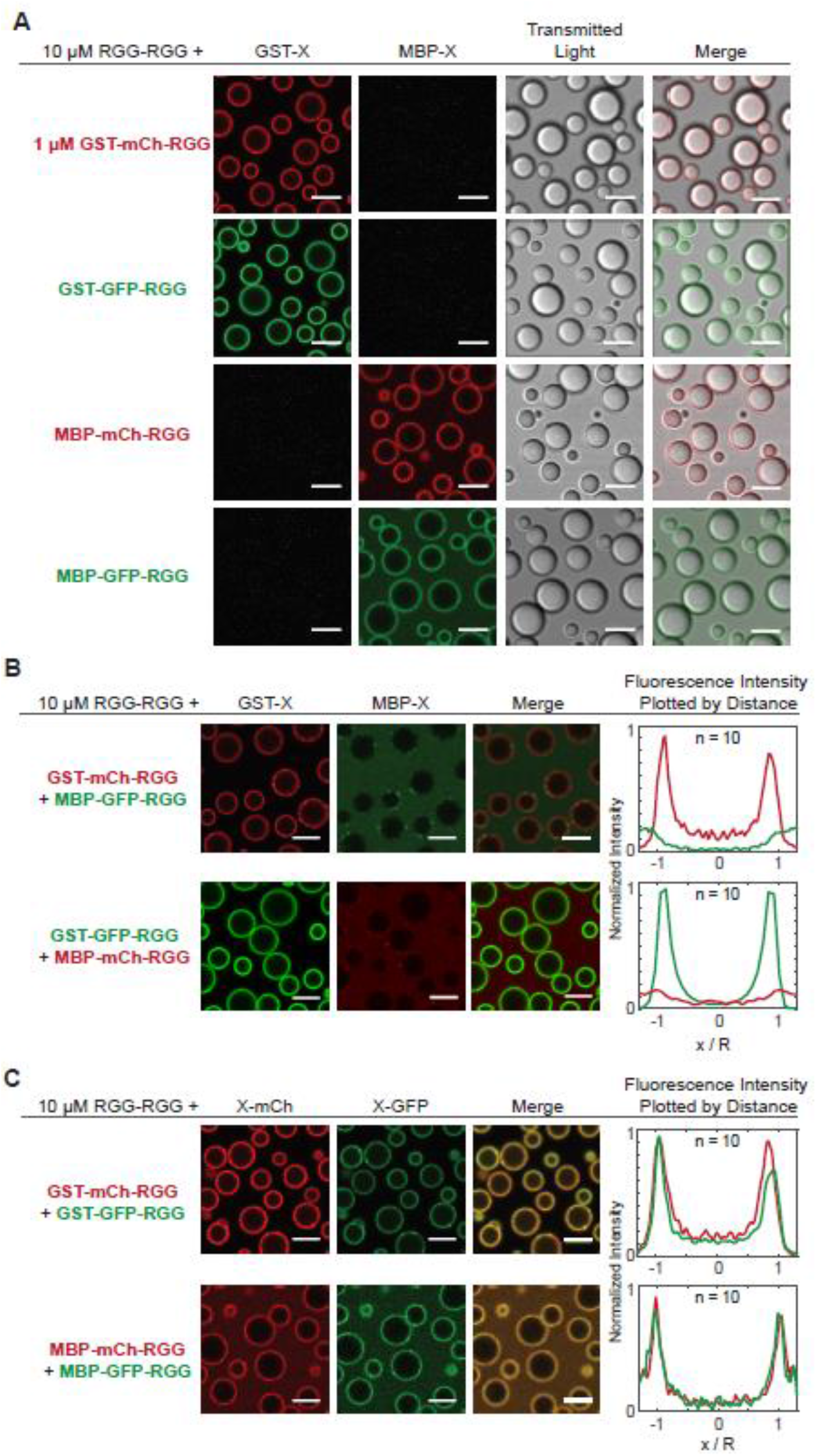
Competitive adsorption to the surface of RGG-RGG condensates when two amphiphilic proteins are introduced. **A**, Films form when a single amphiphilic protein (1 μM) is mixed with RGG-RGG (10 μM), independent of soluble domain (GST or MBP) and fluorophore (mCherry or GFP) combination used. **B**, Combining two dissimilar amphiphilic proteins with RGG-RGG revealed competitive adsorption. Proteins with a GST domain adsorb to the RGG-RGG droplet surface, irrespective of fluorophore included. Amphiphilic proteins with an MBP domain do not adsorb to the surface. Fluorescence intensity line profiles quantified across single condensates confirm GST-containing proteins preferentially adsorb to the surface of the RGG-RGG condensates. n, number of condensates. **C**, Simultaneous adsorption, rather than competition, is observed when combining RGG-RGG with two similar amphiphilic proteins with the same soluble domain (GST or MBP), but different fluorophores. Scale bars, 5 μm. Fluorescence intensity line profiles quantified across single condensates confirm simultaneous adsorption. n, number of condensates.

In agreement with our hypothesis, when we mixed the two classes of surfactant proteins together with RGG-RGG, GST-based proteins displaced MBP-based proteins from the condensate interface (Fig. 5B). When GST-mCherry-RGG and MBP-GFP-RGG together were mixed with RGG-RGG, we observed red rings of GST-mCherry-RGG surrounding RGG-RGG droplets, while MBP-GFP-RGG appeared mostly in the continuous phase. Likewise, when GST-GFP-RGG was mixed with MBP-mCherry-RGG, green rings of GST-GFP-RGG formed while red fluorescence was distributed throughout the continuous phase, indicating MBP-mCherry-RGG was displaced from the interface. Plotting intensity versus radial distance confirmed that proteins with a GST tag remained adsorbed to the surface, whereas MBP-tagged proteins were displaced, irrespective of the fluorescent tag used. As a control, we sought to further verify that the choice of fluorophore has negligible impact on amphiphilic protein adsorption to the RGG-RGG surface (Fig 5C). We mixed RGG-RGG, GST-GFP-RGG, and GST-mCherry-RGG. Separately, we mixed RGG-RGG plus the two MBP-based proteins with differing fluorophores. In both cases, we observed colocalized fluorescent rings in the mCherry and GFP channels via confocal microscopy, resulting in yellow rings in the merged channel view. This suggests that amphiphiles with the same RGG-phobic domain do not displace each other, and are therefore likely to have similar binding affinities to the interface.

Finally, we asked whether this competition phenomenon is concentration dependent. To test this, we prepared RGG-RGG (10 μM) plus MBP-mCherry-RGG (1 μM) with varying GST-GFP-RGG concentrations (SI Appendix Fig. 6). Interestingly, with only 0.1 μM GST-GFP-RGG, MBP-mCherry-RGG remained enriched at the RGG-RGG interface. As we increased the concentration of GST-GFP-RGG, it out-competed MBP-mCherry-RGG for binding, causing decreased red fluorescence at the droplet surface. These results suggest that GST-tagged surfactant proteins displace MBP-tagged surfactant proteins through competitive binding at the interface. This phenomenon may be due to higher solubility of the MBP tag relative to the GST tag. The higher solubility of MBP-based proteins can also be observed in the sample containing MBP-mCherry-RGG and MBP-GFP-RGG, where the continuous phase is more fluorescent as compared to in the system containing GST-mCherry-RGG and GST-GFP-RGG (Fig. 5C). Concomitantly, the strong interactions between the GST domains suggested by its multiphasic behavior (Fig. 4) may also be contributing to localizing the GST-based constructs at the interface.

### Domain Interactions Determine Behavior of Amphiphilic Proteins

We aimed to synthesize a mechanistic explanation for the divergent behaviors we observed with MBP- and GST-based amphiphilic proteins. To summarize these behaviors, GST-based amphiphiles display low partitioning in solution and phase separate at concentrations above 1 μM. MBP-based amphiphiles are relatively more available in solution and do not phase separate at high concentrations, rather controlling condensate size. When mixed together, GST-based amphiphiles outcompete MBP-based amphiphiles for the surface of RGG-RGG condensates. Our analyses also rule out that the fluorophore has a significant impact on amphiphile behavior, indicating that properties intrinsic to GST and MBP are critical. To account for the behaviors observed, we hypothesized that the relative strength of interactions between domains, i.e., homotypic interactions between MBP: MBP and GST: GST, and heterotypic interactions between amphiphilic proteins and the RGG-RGG core, determine both the degree of partitioning of the amphiphile to the surface as well as the formation of hierarchical structures. As mentioned earlier, strong interactions between either GST: RGG or GST: GST could be driving the formation of the multiphasic systems observed.

To explore the role of protein interactions further, we built a computational model that could recapitulate our experimental observations and provide insight into the role of interactions in determining amphiphilic protein behavior. This highly simplified model allowed us to simulate multicomponent phase separation using molecular dynamics (MD) simulations. The computational efficiency of the model allows us to scan a wide parameter space of inter-domain interactions and protein concentrations. Based on our experimental observations, two assumptions were made: (1) the fluorescent tag does not play a major role in dictating the observed behavior and hence can be excluded from this simplified model; (2) among all interactions possible, homotypic interactions between RGG domains are the strongest, supported by its high propensity to phase separate. All other heterotypic and homotypic interaction strengths in the system are scaled with respect to the strength of homotypic RGG interaction.

We first examined a two-component system with RGG-RGG and either MBP-RGG or GST-RGG. To decide on a suitable range of strengths for heterotypic interactions (RGG: GST and RGG: MBP) and homotypic interactions (GST: GST and MBP: MBP), we spanned the parameter space searching for results that approximated our experimental observations. To do so, we calculated the dilute, interfacial, and dense phase concentrations of the molecules across a range of interaction strengths and found those values where molecules are enriched in a similar pattern as observed in our experiments (SI Appendix Figs. 8, 9). Our aim was to identify interaction criteria where GST-RGG and MBP-RGG form a layer surrounding RGG-RGG and upon increasing the concentration of GST-RGG and MBP-RGG, we observe phase separation and accumulation in the dilute phase for the two cases, respectively.

Results from the simulation allow us to make several conclusions regarding the strength of homotypic and heterotopic interactions between the GST, MBP, and RGG domains. First, if RGG: GST or RGG: MBP interaction strengths were high, we would observe partitioning of GST-RGG and MBP-RGG within the RGG-RGG condensates, which is not observed experimentally (SI Appendix Figs. 7C-D, 9). Interaction between the RGG domains in GST-RGG and MBP-RGG is sufficient to enrich those molecules at the surface of RGG-RGG condensates; additional interaction between the folded and phase separating domains shift the nature of the amphiphilic protein towards partitioning within the RGG-RGG core (SI Appendix Figs. 7C-D, 9). Interestingly, this informs our search for the type of interaction that is responsible for driving GST-GFP-RGG phase separation, suggesting that GST: RGG interactions may play less of a role compared to GST: GST interactions in driving GST-GFP-RGG phase separation. Second, if MBP: MBP or GST: GST interaction strength is too low, we lose the formation of a layer of GST-RGG or MBP-RGG surrounding the RGG-RGG core, showing that homotypic interaction between either of the folded domains is important for their enrichment at the RGG-RGG core surface (SI Appendix Figs. 7A-B, 9).

Next, we examined a three component system with RGG-RGG, GST-RGG, and MBP-RGG. Within the parameter space explored, we were able to identify values for the strength of RGG: GST, RGG: MBP, GST: GST and MBP: MBP interactions where the model agrees with our experimental data. We set the GST: GST interaction strength to be greater than or equal to 0.8 times the RGG: RGG interaction strength and the MBP: MBP interaction strength to be less than or equal to 0.2 times that of RGG: RGG interaction strength. Results show RGG-RGG phase separating into droplets (Fig. 6A, left panel), GST-RGG becoming highly enriched at the droplet surface (Fig. 6A, middle panel), while MBP-RGG displays a weaker localization to the interface (Fig. 6A, right panel). Their simulated behaviors in the dilute phase also agrees with that observed experimentally, with GST-RGG being relatively more enriched at the RGG-RGG interface compared to MBP-RGG that is more present in the surrounding solution (Fig. 6B). We can also quantify the distribution of each component relative to the component’s overall concentration (Fig. 6C). RGG-RGG becomes 100 times more concentrated as it phase separates into condensates centralized at r = 0. GST-RGG is highly enriched (~50x) at the interface of RGG-RGG droplets at around r = 7 nm, and thus reducing its relative concentration outside the droplet. In comparison, though we still observe some localization of MBP-RGG at the interface, it mostly remains in the surrounding solution.

**Figure 6:**
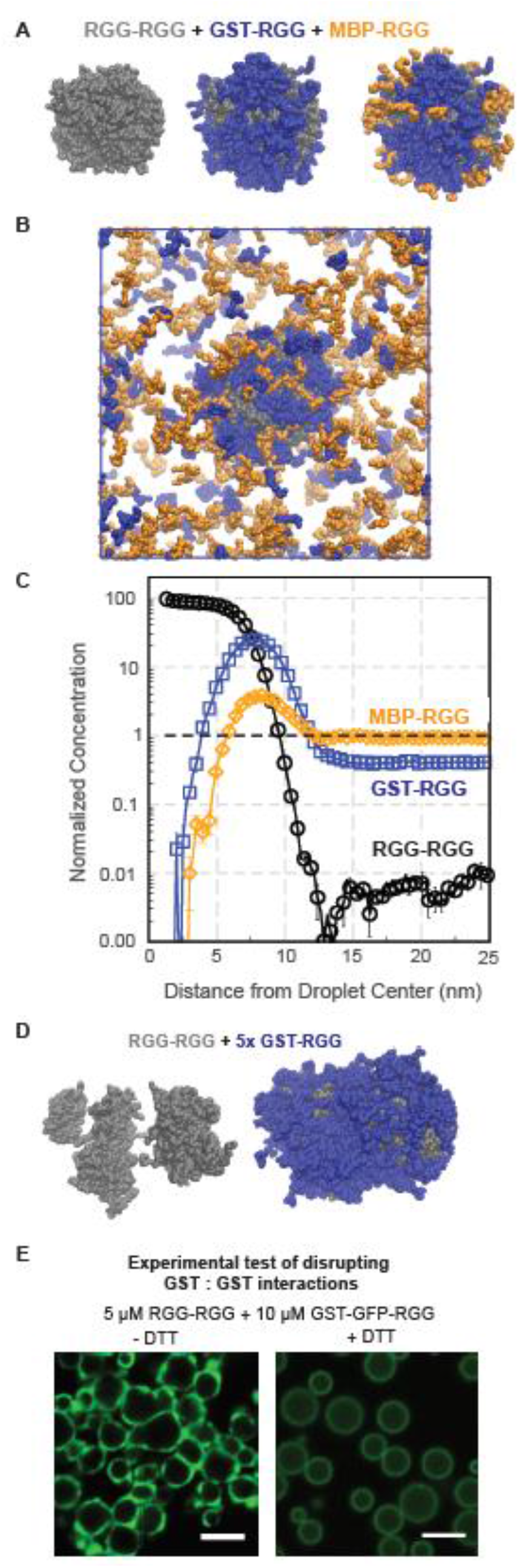
Simulations of RGG-RGG, GST-RGG, and MBP-RGG interactions. **A,** Simulation snapshots of a condensate formed from a ternary mixture with equal concentrations of RGG-RGG (grey), GST-RGG (blue), and MBP-RGG (orange). All images are from the same snapshot, highlighting surface concentrations of different sets of components. GST-RGG displays a higher degree of coverage of the RGG-RGG surface compared to MBP-RGG. **B**, Simulation snapshot displaying the condensed and dilute phases from part A. Higher concentration of MBP-RGG relative to GST-RGG is observed in the dilute phase. **C**, Quantification of protein concentration distribution in space. Protein concentration is normalized to the total concentration of each species in the solution. Mixing MBP-RGG and GST-RGG with RGG-RGG results in greater GST-RGG localization to the surface while MBP-RGG remains in solution. **D**, MD simulation of RGG-RGG with 5x GST-RGG concentration. RGG-RGG forms multiple cores (left) surrounded by a GST-RGG phase (right). **E**, Confocal fluorescence imaging of RGG-RGG (5 μM) and GST-GFP-RGG (10 μM) without DTT (left) and with DTT (5 mM, right). Addition of DTT to RGG-RGG and GST-GFP-RGG mixture results in loss of multiphasic behavior by disruption of GST: GST disulfide bonds. Scale bars, 5 μm.

To build on these observations, we explored whether the interaction values examined in the prior simulation could also produce the multiphasic behavior observed with GST-GFP-RGG. We performed a simulation of a binary mixture of RGG-RGG and GST-RGG, with GST-RGG concentration 5 times higher than used previously. Remarkably, the model predicts GST-RGG phase separation, represented by a dense layer of GST-RGG surrounding the RGG-RGG core. In some scenarios, multiple RGG-RGG cores can be observed within a GST-RGG phase, similar to the experimental observation of a multi-phasic system (Fig. 6D). A reasonable explanation for this behavior is that by splitting the RGG-RGG droplet into smaller clusters, there is an increase in surface area for GST-RGG to interact with RGG-RGG molecules. Furthermore, these results support that strong homotypic interactions between GST domains drive the propensity of GST-based amphiphiles to be highly enriched at the condensate surface and to phase separate at higher concentrations.

Based on this information, we explored potential drivers for the strong GST: GST interaction. It is known that GST has a tendency to dimerize, as well as oligomerize due to 4 exposed cysteine residues at the protein’s surface. The dissociation constant Kd of GST is approximately 1 μM, above which dimerization is preferred.^47^ We hypothesized these homotypic interactions could contribute to the tendency of GST-GFP-RGG to phase separate, as seen in our experimental observations. To test this hypothesis, we assessed the impact of adding a reducing agent that would disrupt the disulfide bonds between GST domains (Fig. 6E). Addition of dithiothreitol (DTT) to a solution of GST-GFP-RGG and RGG-RGG results in the formation of a homogenous film of amphiphilic protein surrounding the RGG-RGG core, as well as increased partitioning in the RGG-RGG phase, irrespective of the concentration of GST-GFP-RGG used. This is in contrast to the partial engulfment and multiphasic behavior observed at high GST-GFP-RGG concentrations in the absence of DTT (Fig. 4, Fig. 6E). This data therefore confirms that GST: GST interactions are necessary for phase separation of the GST-based amphiphiles. Collectively, these molecular simulations and experiments provide mechanistic insight into the differences between the two classes of surfactant proteins tested.

## Discussion

In this paper, we demonstrate that surfactant-like proteins can contribute to the structure and regulation of biomolecular condensates. We report that a two-component in vitro system, consisting of a phase-separating protein and an amphiphilic protein, can reconstitute a range of condensate structures observed in nature. The amphiphilic protein is composed of a domain that is prone to phase separate and a domain that does not phase separate, so at suitable concentrations it adsorbs to the surface of a phase-separated droplet, forming a film. Analogous to surfactant films that stabilize emulsified oil droplets, we discovered that surfactant proteins can stabilize the size of protein condensates. We also found that we can enzymatically disassemble the surfactant film and thereby trigger droplet fusion, demonstrating tunable control of condensate properties by enzymatic manipulation of surfactant proteins. Furthermore, by varying the concentrations of RGG-RGG and the GST-based amphiphiles, a rich phase behavior can be observed, reminiscent of the multiphase and multilayer structures exhibited by naturally occurring condensates. Our results highlight that different amphiphilic proteins can vary in their behavior, one consequence of which is competitive adsorption. Using a simplified model, we show that the strength of interaction between protein domains determine the type of behavior the amphiphilic protein displays. Future studies should further examine the effect on surfactant behavior of mutagenesis, choice of domains, orthogonality of the system and domain architecture for both the core and surfactant proteins.

One outstanding question that our work helps address is how the size of biomolecular condensates is controlled^48,49^. Thermodynamics suggests that smaller droplets should fuse into a single large condensate to reduce the interfacial area between the condensed and dilute phases, thereby minimizing the free energy of the system. However, frequently, cellular condensates coexist as small droplets that do not fuse. Several explanations have been proposed and are likely at play^39,40,57,32^. Our work provides insight into an important alternative mechanism through which droplet size may be regulated. For certain amphiphilic proteins that we tested, presence of the adsorbed surface layer stabilized droplets against fusion. Relatedly, one recent computational study demonstrated that low-valency client molecules that partition with scaffold proteins could theoretically modulate condensate size by acting as surfactants and affecting droplet surface tension^52^. Our findings raises the possibility that cells could be leveraging surfactant-like, amphiphilic proteins to control organelle surface tension and size. To elucidate the mechanism through which size stabilization is occurring in our system, further research quantifying the surface tension of condensates is necessary. There are also future directions in classifying the type of surfactant and emulsion observed in our system, given the important distinction between emulsions and microemulsions in their behavior and applications^53^.

Our results also contribute to the growing body of literature on naturally-occurring, multi-layered condensates. The coexistence of multiple phases or other hierarchical structures has been demonstrated to play important roles in cellular function, such as in the organization of protein-RNA vesicles^54^, TDP-43 droplets^55^, the nucleolus^56^, FMRP/CAPRIN1 droplets^45^, stress granules^57^ and P granules^14^. These hierarchical structures have been described using various terminologies, including core-shell structures^12^, liquid spherical shells^55^, and Pickering emulsions^58^. These structures have been associated with several roles, including catalytic shells^12^ and selective barriers^55^. Beyond the field of liquid-liquid phase separation of IDPs, the film morphologies we observe also resemble naturally occurring plasma membrane vesicles^59^ as well as apolipoprotein micelles^60^. Hence, multiphasic or multilayer morphologies can display a wide range of behaviors and play a variety of roles in cells. Our engineered system of two proteins is reminiscent of several of these behaviors and assemblies and can therefore be useful for understanding the principles of formation of multi-layered condensates. A potential future direction is to conduct a bioinformatic search for naturally-occurring intracellular proteins with amphiphilic properties and systematically test whether these proteins, if any, partition to the surface of known membraneless organelles.

Our system is also similar to other synthetic systems and may also serve as a base platform for designing new bio-inspired materials. Related to our work, formation of spherical micelles have been observed with elastin-like polypeptides (ELPs)^61^, amphiphilic polymers^62,63,64^ as well as plasma proteins forming core-shell structures at the air-water interface^65^. Furthermore, there is a growing field concerned with developing condensates with controllable material properties with applications in several areas such as therapeutic delivery, metabolic engineering, and chemical separations^66–68^. We demonstrate that we can develop a multi-phase system with controllable surface patterning depending on the construction of a modular surfactant protein. Future investigations could aim at imparting function onto our film system such as by engineering two-step reactions.

Membrane-forming lipids are the prototypical example of biological amphiphiles and are essential for the function of membrane-bound compartments^69^. But what role do amphiphilic proteins play in the formation and function of membrane-less compartments? Our work advances the concept that amphiphilic proteins may contribute to the assembly and size regulation of membrane-less organelles. Simple systems comprising phase-separating and amphiphilic proteins, such as those described here, can be further leveraged as models for elucidating fundamental principles governing self-assembly of film and multiphasic hierarchical condensates. This work aims to serve as a basis for investigating the potentially widespread role of amphiphilic proteins in biology.

## Methods

### Cloning

All genes of interest were cloned into pET vectors in frame with C-terminal 6x-His tags. RGG-RGG was cloned as previously described^33^. Other constructs were cloned by PCR and DNA assembly (NEBuilder HiFi DNA Assembly Master Mix; New England Biolabs). The RGG domain used here is the N-terminal IDR (residues 1-168) of C. elegans P granule protein LAF-1^34^. Gene sequences were verified by Sanger sequencing (Genewiz).

### Protein expression and purification

For bacterial expression, plasmids were transformed into BL21(DE3) competent *E. coli* (New England BioLabs). Colonies picked from fresh plates were grown for 8h at 37 °C in 1 mL LB + 1% glucose while shaking at 250 rpm. This starter culture (0.5 mL) was then used to inoculate 0.5 L cultures. For surfactant proteins (GST- and MBP-based constructs), cultures were grown overnight in 2 L baffled flasks in Terrific Broth medium (Fisher Scientific) supplemented with 4 g/L glycerol at 18°C while shaking at 250 rpm. Once the OD600 reached approximately 1, expression was induced with 500 μM isopropyl β-D-1-thiogalactopyranoside (IPTG). For RGG-RGG, cultures were grown in 2 L baffled flasks in Autoinduction medium (Formedium) supplemented with 4 g/L glycerol at 37 °C overnight while shaking at 250 rpm. The pET vectors used contained a kanamycin resistance gene; kanamycin was used at concentrations of 50 μg/mL in cultures^70^. After overnight expression at 18 °C or 37 °C, bacterial cells were pelleted by centrifugation at 4100 x g at 4 °C. Pellets were resuspended in lysis buffer (1 M NaCl, 20 mM Tris, 20 mM imidazole, Roche EDTA-free protease inhibitor, pH 7.5) and lysed by sonication. Lysate was clarified by centrifugation at 25000 x g for 30 minutes at 25 °C. The clarified lysate was then filtered with a 0.22 μm filter. Lysis was conducted on ice, but other steps were conducted at room temperature to prevent phase separation.

Proteins were purified using an AKTA Pure FPLC with 1 mL nickel-charged HisTrap columns (Cytiva) for affinity chromatography of the His-tagged proteins. After injecting proteins onto the column, the column was washed with 500 mM NaCl, 20 mM Tris, 20 mM imidazole, pH 7.5. Proteins were eluted with a linear gradient up to 500 mM NaCl, 20 mM Tris, 500 mM imidazole, pH 7.5. Proteins were dialyzed overnight using 7 kDa MWCO membranes (Slide-A-Lyzer G2, Thermo Fisher) into physiological buffer (150 mM NaCl, 20 mM Tris, pH 7.5), with the exception that GST-GFP-RGG was dialyzed into high salt buffer (1M NaCl, 20mM Tris, pH 7.5). RGG-RGG was dialyzed at 45°C to inhibit phase separation because phase-separated protein bound irreversibly to the dialysis membrane; other proteins were dialyzed at room temperature.

Proteins were snap frozen in liquid N2 in single-use aliquots and stored at −80 °C. For microscopy experiments, protein samples were prepared as follows: RGG-RGG protein aliquots were thawed above the phase transition temperature. MBP- and GST-based proteins were thawed at room temperature. GST-GFP-RGG protein aliquots were concentrated with a centrifugal concentrator (Amicon Ultra Centrifugal Filters, Millipore). Proteins were then mixed with 20 mM Tris, 150 mM NaCl, pH 7.5 with the exception of GST-GFP-RGG which was mixed with a combination of 20 mM Tris, 150 mM NaCl, pH 7.5 and 20 mM Tris, 0 mM NaCl, pH 7.5 buffers to obtain a final solution containing 150 mM NaCl. All experiments were conducted with a buffer concentration of 20 mM Tris, 150 mM NaCl, pH 7.5. Protein concentrations were measured based on their absorbance at 280 nm using a Nanodrop spectrophotometer (ThermoFisher); RGG-RGG and GST-GFP-RGG were mixed in a 1:1 ratio with 8 M urea to prevent phase separation during concentration measurements.

### SDS-PAGE

For chromatographically purified proteins, SDS-PAGE was run using NuPAGE 4-12% Bis-Tris gels (Invitrogen) and stained using a Coomassie stain (SimplyBlue SafeStain; Invitrogen).

### Microscopy: phase behavior, TEV, and FRAP

RGG-RGG was mixed with MBP-based and/or GST-based amphiphiles at the desired protein concentrations while maintaining the buffer conditions at 150 mM NaCl, 20 mM Tris, pH 7.5. Protein samples were plated on 16-well glass-bottom dishes (#1.5 glass thickness; Grace Bio-Labs) that were coated with 5% Pluronic F-127 (Sigma-Aldrich) for a minimum of 10 minutes. The chambers were washed with buffer solution prior to plating the protein samples.

Imaging was performed on a Zeiss Axio Observer 7 inverted microscope equipped with an LSM900 laser scanning confocal module and employing a 63x/1.4 NA plan-apochromatic, oil-immersion objective. GFP was excited to fluoresce with a 488 nm laser, and mCherry was excited with a 561 nm laser. Confocal fluorescence images were captured using GaAsP detectors. Transmitted light images were collected with either the ESID module or an Axiocam 702 sCMOS camera (Zeiss), in both cases using a 0.55 NA condenser.

Time-resolved experiments with TEV protease were performed by adding 1 μL TEV protease (New England Biolabs) after the 3rd time point to an imaging well containing 100 μL protein sample. Size dependence experiments were performed by mixing RGG-RGG with 1 or 5 μM MBP-RGG-GFP and imaged after 1 hour of equilibration. Experiments with dithiothreitol (DTT) were done by mixing it with protein sample at 5mM. All microscopy experiments were conducted at 14-17 °C.

### Droplet image analysis

Image analysis and data processing were performed in MATLAB R2020b. The fluorescence intensity profile of the condensates with fluorescent-tagged proteins was measured by using the Hough Transform to identify droplet locations and drawing a line that spanned the droplet diameter plus 1/4th of a radius length in each direction across the droplets. Line-scan graphs were generated in MATLAB. The number of analyzed condensates (n) is indicated in each figure. The total fluorescence intensity of the shell region of the condensates was measured by manually tracking fluorescence over a background threshold across the z-stack over time. Total intensity graphs were generated in MATLAB. Condensate size was calculated using MATLAB’s inbuilt Hough Transform function. Box and whisker plot and scatter plots were generated in Excel. Line graphs were generated in MATLAB.

### Molecular dynamics simulations

We represent the individual protein sequences i.e., RGG (A), GST (B) and MBP (C) as 20 monomer long disordered domains with identical harmonic bonds connecting all monomers in a domain and also connecting domains with each other to form tandem A-A, A-B and A-C chains. The total interaction energy in our simplistic model includes contributions from this harmonic bond potential and short-range van der Waals (vdW) interactions between monomers,

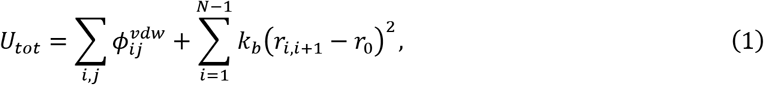

where *k_b_* is the spring constant = 20 kJ/Å^2^, and r_0_ is the equilibrium bond length = 3.81 Å in the harmonic bond potential, and N = 40 is the total number of beads in a tandem chain. The interactions between monomers i and j are represented using the following Ashbaugh-Hatch functional form with σ_*ij*_ = 5 Å and ϵ= 0.2 kcal/mol for all heterotypic and homotypic pairs,

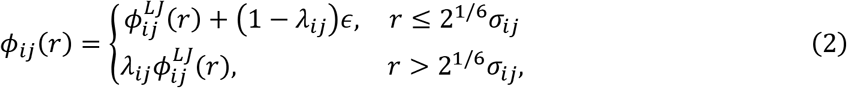

where 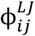 is the standard Lennard-Jones (LJ) potential shown below:

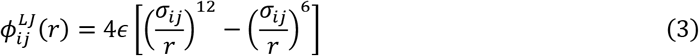

Here *λ_ij_* controls the interaction strength between monomers i and j.

All simulations were conducted at 298K with 250 chains of A-A while the number of X-A (X = B or C) chains were varied, in a cubic periodic box of size 50 nm. The simulations were conducted using HOOMD-Blue and coexistence densities were calculated using the protocol mentioned in our previous work^71^.

## Data Availability Statement

All the quantitative analyses discussed in this paper were generated based on computer codes that we will make publicly available (links will be made available upon publication).

## Acknowledgements

We acknowledge Zheng Shi, Dragomir Milovanovic, Christian Hoffmann, Julie Forman-Kay, Omar Adame-Arana, and Samuel Safran for valuable feedback. BSS thanks Daniel Hammer and Matthew Good for guidance during early stages of this project. We also thank Xinyi Li for her assistance with protein purification. This research was supported by Rutgers University startup funds and a Research Council Award (to BSS). FMK is supported by National Institutes of Health (NIH) grant T32 GM135141. RMR and JM are supported by NIH grant R01 GM120537 (to JM). Use of the high-performance computing capabilities of the Extreme Science and Engineering Discovery Environment (XSEDE), which is supported by NSF grant TG-MCB-120014, is gratefully acknowledged.

## Author Contributions

FMK, BF, and BSS conceived and designed research. FMK, BF, and BSS conducted wet lab experiments. RMR and JM designed the computer experiments. RMR conducted the molecular simulations. BSS and JM supervised research. FMK, BF, RMR, JM, and BSS wrote manuscript.

## References

1. Wilson, E. B. The Structure of the Protoplasm. Science 10, 33–45 (1899).

2. Banani, S. F., Lee, H. O., Hyman, A. A. & Rosen, M. K. Biomolecular condensates: Organizers of cellular biochemistry. Nature Reviews Molecular Cell Biology vol. 18 285–298 (2017).

3. Shin, Y. & Brangwynne, C. P. Liquid phase condensation in cell physiology and disease. Science 357, eaaf4382 (2017).

4. Lyon, A. S., Peeples, W. B. & Rosen, M. K. A framework for understanding the functions of biomolecular condensates across scales. Nature Reviews Molecular Cell Biology vol. 22 215–235 (2021).

5. Alberti, S. & Dormann, D. Liquid-Liquid Phase Separation in Disease. Annu. Rev. Genet. 53, 171–194 (2019).

6. Schuster, B. S. et al. Biomolecular Condensates: Sequence Determinants of Phase Separation, Microstructural Organization, Enzymatic Activity, and Material Properties. Journal of Physical Chemistry B (2021) doi:10.1021/acs.jpcb.0c11606.

7. Hawgood, S. & Clements, J. A. Pulmonary surfactant and its apoproteins. Journal of Clinical Investigation vol. 86 1–6 (1990).

8. Veldhuizen, E. J. A. & Haagsman, H. P. Role of pulmonary surfactant components in surface film formation and dynamics. Biochimica et Biophysica Acta - Biomembranes vol. 1467 255–270 (2000).

9. Hyman, A. A., Weber, C. A. & Jülicher, F. Liquid-Liquid Phase Separation in Biology. Annu. Rev. Cell Dev. Biol. 30, 39–58 (2014).

10. Cuylen, S. et al. Ki-67 acts as a biological surfactant to disperse mitotic chromosomes. Nature 535, 308–312 (2016).

11. Yamasaki, A. et al. Liquidity Is a Critical Determinant for Selective Autophagy of Protein Condensates. Mol. Cell 77, 1163–1175.e9 (2020).

12. Gallego, L. D. et al. Phase separation directs ubiquitination of gene-body nucleosomes. Nature 579, 592–597 (2020).

13. Feric, M. et al. Coexisting Liquid Phases Underlie Nucleolar Subcompartments. Cell 165, 1686–1697 (2016).

14. Putnam, A., Cassani, M., Smith, J. & Seydoux, G. A gel phase promotes condensation of liquid P granules in Caenorhabditis elegans embryos. Nat. Struct. Mol. Biol. 26, 220–226 (2019).

15. Jain, S. et al. ATPase-Modulated Stress Granules Contain a Diverse Proteome and Substructure. Cell 164, 487–498 (2016).

16. Fei, J. et al. Quantitative analysis of multilayer organization of proteins and RNA in nuclear speckles at super resolution. J. Cell Sci. 130, 4180–4192 (2017).

17. West, J. A. et al. Structural, super-resolution microscopy analysis of paraspeckle nuclear body organization. J. Cell Biol. 214, (2016).

18. Boeynaems, S. et al. Spontaneous driving forces give rise to protein-RNA condensates with coexisting phases and complex material properties. Proc. Natl. Acad. Sci. U. S. A. (2019) doi:10.1073/pnas.1821038116.

19. Fisher, R. S. & Elbaum-Garfinkle, S. Tunable multiphase dynamics of arginine and lysine liquid condensates. Nat. Commun. 11, 4628 (2020).

20. Kaur, T. et al. Sequence-encoded and composition-dependent protein-RNA interactions control multiphasic condensate morphologies. Nat. Commun. 12, 2020.08.30.273748 (2021).

21. Swain, P. & Weber, S. C. Dissecting the complexity of biomolecular condensates. Biochem. Soc. Trans. 48, 2591–2602 (2020).

22. Sanders, D. W. et al. Competing Protein-RNA Interaction Networks Control Multiphase Intracellular Organization. Cell 181, 306–324.e28 (2020).

23. Wan, G. et al. Spatiotemporal regulation of liquid-like condensates in epigenetic inheritance. Nature 557, 679–683 (2018).

24. Israelachvili, J. Intermolecular and Surface Forces. Intermolecular and Surface Forces (Elsevier Inc., 2011). doi:10.1016/C2009-0-21560-1.

25. Brangwynne, C. P. Phase transitions and size scaling of membrane-less organelles. J. Cell Biol. 203, 875–81 (2013).

26. Klein, I. A. et al. Partitioning of cancer therapeutics in nuclear condensates. Science (80-.). 368, 1386–1392 (2020).

27. Welsh, T. et al. Single particle zeta-potential measurements reveal the role of electrostatics in protein condensate stability. bioRxiv 2020.04.20.047910 (2020) doi:10.1101/2020.04.20.047910.

28. Feric, M. & Brangwynne, C. P. A nuclear F-actin scaffold stabilizes ribonucleoprotein droplets against gravity in large cells. Nat. Cell Biol. 15, 1253–1259 (2013).

29. Quiroz, F. G. et al. Liquid-liquid phase separation drives skin barrier formation. Science 367, (2020).

30. Snead, W. T., Gerbich, T. M., Seim, I., Hu, Z. & Gladfelter, A. S. Membrane surfaces regulate assembly of a ribonucleoprotein condensate. bioRxiv 2021.04.24.441251 (2021) doi:10.1101/2021.04.24.441251.

31. Ranganathan, S. & Shakhnovich, E. I. Dynamic metastable long-living droplets formed by sticker-spacer proteins. Elife 9, 1–25 (2020).

32. Wurtz, J. D. & Lee, C. F. Chemical-Reaction-Controlled Phase Separated Drops: Formation, Size Selection, and Coarsening. Phys. Rev. Lett. 120, 078102 (2018).

33. Schuster, B. S. et al. Controllable protein phase separation and modular recruitment to form responsive membraneless organelles. Nat. Commun. 9, 1–12 (2018).

34. Elbaum-Garfinkle, S. et al. The disordered P granule protein LAF-1 drives phase separation into droplets with tunable viscosity and dynamics. Proc. Natl. Acad. Sci. U. S. A. 112, 7189–7194 (2015).

35. Schuster, B. S. et al. Identifying sequence perturbations to an intrinsically disordered protein that determine its phase-separation behavior. Proc. Natl. Acad. Sci. U. S. A. 117, (2020).

36. Chong, P. A., Vernon, R. M. & Forman-Kay, J. D. RGG/RG Motif Regions in RNA Binding and Phase Separation. J. Mol. Biol. 430, 4650–4665 (2018).

37. Esposito, D. & Chatterjee, D. K. Enhancement of soluble protein expression through the use of fusion tags. Curr. Opin. Biotechnol. 17, 353–358 (2006).

38. Kapust, R. B. & Waugh, D. S. Escherichia coli maltose-binding protein is uncommonly effective at promoting the solubility of polypeptides to which it is fused. Protein Sci. 8, 1668–1674 (1999).

39. Reed, E. H., Schuster, B. S., Good, M. C. & Hammer, D. A. SPLIT: Stable Protein Coacervation Using a Light Induced Transition. ACS Synth. Biol. 9, 500–507 (2020).

40. Caldwell, R. M. et al. Optochemical Control of Protein Localization and Activity within Cell-like Compartments. Biochemistry 57, 2590–2596 (2018).

41. Kapust, R. B. et al. Tobacco etch virus protease: Mechanism of autolysis and rational design of stable mutants with wild-type catalytic proficiency. Protein Eng. 14, 993–1000 (2001).

42. Tcholakova, S., Denkov, N. D. & Banner, T. Role of surfactant type and concentration for the mean drop size during emulsification in turbulent flow. Langmuir 20, 7444–7458 (2004).

43. Wang, F., Richards, V. N., Shields, S. P. & Buhro, W. E. Kinetics and mechanisms of aggregative nanocrystal growth. Chem. Mater. 26, 5–21 (2014).

44. Linsenmeier, M., Kopp, M. R. G., Stavrakis, S., de Mello, A. & Arosio, P. Analysis of biomolecular condensates and protein phase separation with microfluidic technology. Biochim. Biophys. Acta - Mol. Cell Res. 1868, 118823 (2021).

45. Kim, T. H. et al. Phospho-dependent phase separation of FMRP and CAPRIN1 recapitulates regulation of translation and deadenylation. Science (80-.). 365, 825–829 (2019).

46. Lu, T. & Spruijt, E. Multiphase Complex Coacervate Droplets. J. Am. Chem. Soc. 142, 2905–2914 (2020).

47. Huang, Y. C., Misquitta, S., Blond, S. Y., Adams, E. & Colman, R. F. Catalytically active monomer of glutathione S-transferase π and key residues involved in the electrostatic interaction between subunits. J. Biol. Chem. 283, 32880–32888 (2008).

48. Weber, S. C. & Brangwynne, C. P. Inverse size scaling of the nucleolus by a concentration-dependent phase transition. Curr. Biol. 25, 641–646 (2015).

49. Dar, F. & Pappu, R. Restricting the sizes of condensates. eLife vol. 9 1–3 (2020).

50. Brangwynne, C. P. et al. Germline P granules are liquid droplets that localize by controlled dissolution/condensation. Science (80-.). 324, 1729–1732 (2009).

51. Berry, J., Brangwynne, C. P. & Haataja, M. Physical principles of intracellular organization via active and passive phase transitions. Reports on Progress in Physics vol. 81 (2018).

52. Sanchez-burgos, I., Joseph, J. A., Collepardo-guevara, R. & Espinosa, J. R. Size conservation emerges spontaneously in biomolecular condensates formed by scaffolds and surfactant clients. (2021) doi:https://doi.org/10.1101/2021.04.30.442154.

53. Andelman, D., Cates, M. E., Roux, D. & Safran, S. A. Structure and phase equilibria of microemulsions. J. Chem. Phys. 87, 7229–7241 (1987).

54. Alshareedah, I., Moosa, M. M., Raju, M., Potoyan, D. A. & Banerjee, P. R. Phase transition of RNA-protein complexes into ordered hollow condensates. Proc. Natl. Acad. Sci. U. S. A. 117, 15650–15658 (2020).

55. Yu, H. et al. HSP70 chaperones RNA-free TDP-43 into anisotropic intranuclear liquid spherical shells. Science (80-.). 371, (2021).

56. Feric, M. et al. Coexisting Liquid Phases Underlie Nucleolar Subcompartments Article Coexisting Liquid Phases Underlie Nucleolar Subcompartments. Cell 165, 1686–1697 (2016).

57. Khong, A. et al. The Stress Granule Transcriptome Reveals Principles of mRNA Accumulation in Stress Granules. Mol. Cell 68, 808–820.e5 (2017).

58. Crowe, C. D. & Keating, C. D. Liquid–liquid phase separation in artificial cells. Interface Focus vol. 8 (2018).

59. Baumgart, T. et al. Large-scale fluid/fluid phase separation of proteins and lipids in giant plasma membrane vesicles. Proc. Natl. Acad. Sci. U. S. A. 104, 3165–3170 (2007).

60. Mahley, R. W., Innerarity, T. L., Rall, S. C. & Weisgraber, K. H. Plasma lipoproteins: Apolipoprotein structure and function. J. Lipid Res. 25, 1277–1294 (1984).

61. Simon, J. R., Carroll, N. J., Rubinstein, M., Chilkoti, A. & López, G. P. Programming molecular self-assembly of intrinsically disordered proteins containing sequences of low complexity. Nat. Chem. 9, 509–515 (2017).

62. Ianiro, A. et al. Controlling the Spatial Distribution of Solubilized Compounds within Copolymer Micelles. Langmuir 35, 4776–4786 (2019).

63. Jiménez-Ángeles, F. et al. Self-Assembly of Charge-Containing Copolymers at the Liquid-Liquid Interface. ACS Cent. Sci. 5, 688–699 (2019).

64. Jiménez-Ángeles, F. et al. Self-Assembly of Charge-Containing Copolymers at the Liquid-Liquid Interface. ACS Cent. Sci. 5, 688–699 (2019).

65. Liao, Z., Lampe, J. W., Ayyaswamy, P. S., Eckmann, D. M. & Dmochowski, I. J. Protein assembly at the air-water interface studied by fluorescence microscopy. Langmuir 27, 12775–12781 (2011).

66. Amiram, M., Luginbuhl, K. M., Li, X., Feinglos, M. N. & Chilkoti, A. Injectable protease-operated depots of glucagon-like peptide-1 provide extended and tunable glucose control. Proc. Natl. Acad. Sci. U. S. A. 110, 2792–2797 (2013).

67. Avalos, J. L., Fink, G. R. & Stephanopoulos, G. Compartmentalization of metabolic pathways in yeast mitochondria improves the production of branched-chain alcohols. Nat. Biotechnol. 31, 335–341 (2013).

68. Roberts, S. et al. Complex microparticle architectures from stimuli-responsive intrinsically disordered proteins. Nat. Commun. 11, 1–10 (2020).

69. Zeno, W. F., Day, K. J., Gordon, V. D. & Stachowiak, J. C. Principles and Applications of Biological Membrane Organization. Annual Review of Biophysics vol. 49 19–39 (2020).

70. Studier, F. W. Stable Expression Clones and Auto-Induction for Protein Production in E. coli. in Methods in molecular biology (Clifton, N.J.) vol. 1091 17–32 (2014).

71. Dignon, G. L., Zheng, W., Kim, Y. C., Best, R. B. & Mittal, J. Sequence determinants of protein phase behavior from a coarse-grained model. PLoS Comput. Biol. 14, 1–23 (2018).

