## Supplemental Materials for "Amphiphilic proteins coassemble into multiphasic condensates and act as biomolecular surfactants"

**This PDF file includes:**

Supplementary Text
Figures S1 to S9
SI References

### 1. Supplementary Text

##### 1.1 Protein sequences used in this work

Throughout this paper, RGG denotes the RGG domain from LAF-1 (residues 1-168 of LAF-1). All proteins included a hexahistidine tag at the C-terminus for immobilized metal affinity chromatography. Below, RGG, GST, MBP, and GFP domains are color-coded; tags, linkers, and cut sites are not colored.

**RGG-RGG**

MESNQSNNGGSGNAALNRGGRYVPPHLRGGDGGAAAAASAGGDDRRGGAGGGGYRRGGGNSGGG
GGGGYDRGYNDNRDDRDNRGGSGGYGRDRNYEDRGYNGGGGGGGNRGYNNNRGGGGGGYNRQD
RGDGGSSNFSRGGYNNRDEGSDNRGSGRSYNNDRRDNGGDGEFGKLMESNQSNNGGSGNAALNRG
GRYVPPHLRGGDGGAAAAASAGGDDRRGGAGGGGYRRGGGNSGGGGGGGYDRGYNDNRDDRDNR
GGSGGYGRDRNYEDRGYNGGGGGGGNRGYNNNRGGGGGGYNRQDRGDGGSSNFSRGGYNNRDE
GSDNRGSGRSYNNDRRDNGGDGLEHHHHHH

**MBP-GFP-RGG**

MKIEEGKLVIWINGDKGYNGLAEVGKKFEKDTGIKVTVEHPDKLEEKFPQVAATGDGPDIIFWAHDRFGG
YAQSGLLAEITPDKAFQDKLYPFTWDAVRYNGKLIAYPIAVEALSLIYNKDLLPNPPKTWEEIPALDKELKA
KGKSALMFNLQEPYFTWPLIAADGGYAFKYENGKYDIKDVGVDNAGAKAGLTFLVDLIKNKHMNADTDYS
IAEAAFNKGETAMTINGPWAWSNIDTSKVNYGVTVLPTFKGQPSKPFVGVLSAGINAASPNKELAKEFLEN
YLLTDEGLEAVNKDKPLGAVALKSYEEELVKDPRIAATMENAQKGEIMPNIPQMSAFWYAVRTAVINAAS
GRQTVDEALKDAQTNSSSNNNNNNNNNNLGETVRFQSMVSKGEELFTGVVPILVELDGDVNGHKFSVS
GEGEGDATYGKLTLKFICTTGKLPVPWPTLVTTLTYGVQCFSRYPDHMKQHDFFKSAMPEGYVQERTIFF
KDDGNYKTRAEVKFEGDTLVNRIELKGIDFKEDGNILGHKLEYNYNSHNVYIMADKQKNGIKVNFKIRHNIE
DGSVQLADHYQQNTPIGDGPVLLPDNHYLSTQSKLSKDPNEKRDHMVLLEFVTAAGITLGMDELYKGGG
SENLYFQGEFGKLMESNQSNNGGSGNAALNRGGRYVPPHLRGGDGGAAAAASAGGDDRRGGAGGGG
YRRGGGNSGGGGGGGYDRGYNDNRDDRDNRGGSGGYGRDRNYEDRGYNGGGGGGGNRGYNNNR
GGGGGGYNRQDRGDGGSSNFSRGGYNNRDEGSDNRGSGRSYNNDRRDNGGDGLEHHHHHH

**MBP-RGG-GFP**

MKIEEGKLVIWINGDKGYNGLAEVGKKFEKDTGIKVTVEHPDKLEEKFPQVAATGDGPDIIFWAHDRFGG
YAQSGLLAEITPDKAFQDKLYPFTWDAVRYNGKLIAYPIAVEALSLIYNKDLLPNPPKTWEEIPALDKELKA
KGKSALMFNLQEPYFTWPLIAADGGYAFKYENGKYDIKDVGVDNAGAKAGLTFLVDLIKNKHMNADTDYS
IAEAAFNKGETAMTINGPWAWSNIDTSKVNYGVTVLPTFKGQPSKPFVGVLSAGINAASPNKELAKEFLEN
YLLTDEGLEAVNKDKPLGAVALKSYEEELVKDPRIAATMENAQKGEIMPNIPQMSAFWYAVRTAVINAAS
GRQTVDEALKDAQTNSSSNNNNNNNNNNLGENLYFQGGMESNQSNNGGSGNAALNRGGRYVPPHLR
GGDGGAAAAASAGGDDRRGGAGGGGYRRGGGNSGGGGGGGYDRGYNDNRDDRDNRGGSGGYGR
DRNYEDRGYNGGGGGGGNRGYNNNRGGGGGGYNRQDRGDGGSSNFSRGGYNNRDEGSDNRGSGR
SYNNDRRDNGGDGEFVSKGEELFTGVVPILVELDGDVNGHKFSVSGEGEGDATYGKLTLKFICTTGKLP
VPWPTLVTTLTYGVQCFSRYPDHMKQHDFFKSAMPEGYVQERTIFFKDDGNYKTRAEVKFEGDTLVNRI
ELKGIDFKEDGNILGHKLEYNYNSHNVYIMADKQKNGIKVNFKIRHNIEDGSVQLADHYQQNTPIGDGPVL
LPDNHYLSTQSKLSKDPNEKRDHMVLLEFVTAAGITLGMDELYKLEHHHHHH

**GST-GFP-RGG**

MSPILGYWKIKGLVQPTRLLLEYLEEKYEEHLYERDEGDKWRNKKFELGLEFPNLPYYIDGDVKLTQSMAI
IRYIADKHNMLGGCPKERAEISMLEGAVLDIRYGVSRIAYSKDFETLKVDFLSKLPEMLKMFEDRLCHKTYL
NGDHVTHPDFMLYDALDVVLYMDPMCLDAFPKLVCFKKRIEAIPQIDKYLKSSKYIAWPLQGWQATFGGG
DHPPGSETVRFQSMVSKGEELFTGVVPILVELDGDVNGHKFSVSGEGEGDATYGKLTLKFICTTGKLPVP
WPTLVTTLTYGVQCFSRYPDHMKQHDFFKSAMPEGYVQERTIFFKDDGNYKTRAEVKFEGDTLVNRIEL
KGIDFKEDGNILGHKLEYNYNSHNVYIMADKQKNGIKVNFKIRHNIEDGSVQLADHYQQNTPIGDGPVLLP
DNHYLSTQSKLSKDPNEKRDHMVLLEFVTAAGITLGMDELYKGGGSENLYFQGEFGKLMESNQSNNGG

SGNAALNRGGRYVPPHLRGGDGGAAAAASAGGDDRRGGAGGGGYRRGGGNSGGGGGGGYDRGYN
DNRDDRDNRGGSGGYGRDRNYEDRGYNGGGGGGGNRGYNNNRGGGGGGYNRQDRGDGGSSNFS
RGGYNNRDEGSDNRGSGRSYNNDRRDNGGDGLEHHHHHH

#### 2. Supplementary Figures

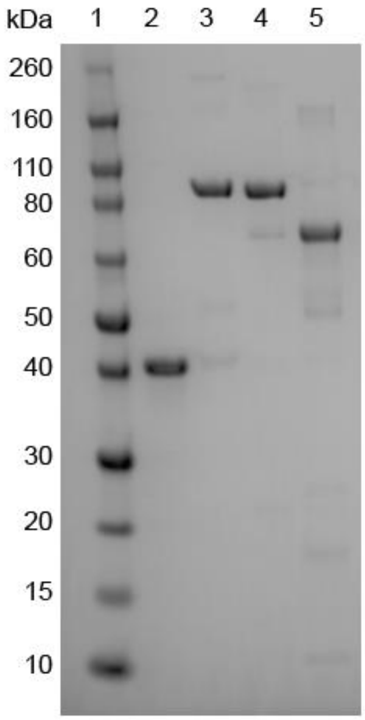


1. Ladder
2. RGG-RGG
3. MBP-GFP-RGG
4. MBP-RGG-GFP
5. GST-GFP-RGG

**Figure S1: SDS-PAGE of purified fusion proteins developed for this study.**

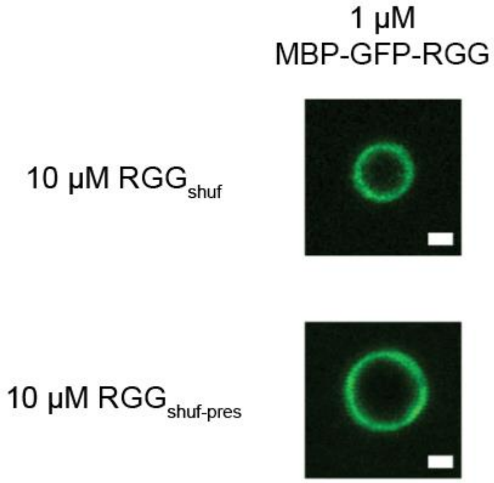


**Figure S2: Formation of surfactant protein film is still observed in a system with RGG variants.** A film of MBP-GFP-RGG can be observed surrounding the core protein droplets in a system that replaces RGG-RGG as the core protein with RGG variants. The variants tested here have the same amino acid composition as wild-type RGG, but the amino acids were shuffled to segregate charges to opposite termini.[1] In one variant ($RGG_{shuf\text{-}pres}$, bottom) the amino acid sequence was shuffled in a manner that preserved the eIF4E-binding motif (located at residues 21–28), whereas the other shuffled variant did not preserve this motif ($RGG_{shuf}$, top).[1] Scale bars, 1 µm.

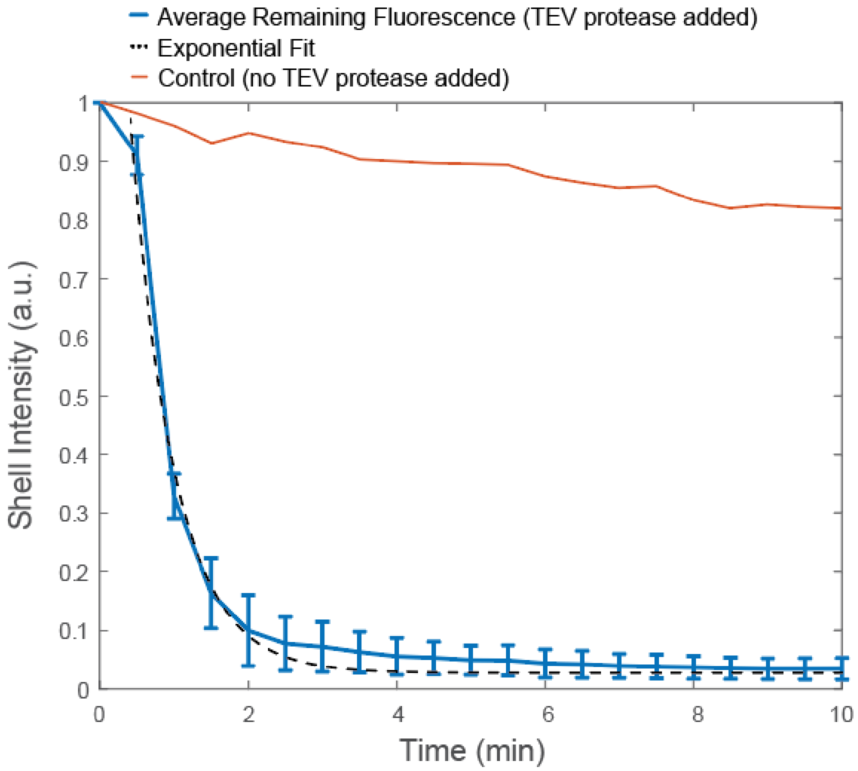


**Figure S3: Control for Figure 2D shows minimal photobleaching occurring.** The experimental data from Fig. 2D is replotted here (blue), showing exponential decay in fluorescence intensity of the surfactant ring when a sample of 10 μM RGG-RGG + 1 μM MBP-GFP-RGG is treated with TEV protease, which cleaves C-terminal to the GFP. Control data (red) represents fluorescence intensity of the surfactant ring from imaging a sample of 10 μM RGG-RGG + 1 μM MBP-GFP-RGG under the same experimental conditions as shown in Fig. 2D but omitting addition of TEV protease. Results show only modest and linear loss of fluorescence intensity for the control, likely due to photobleaching, as compared to the exponential and substantial loss of fluorescence intensity with TEV protease addition.

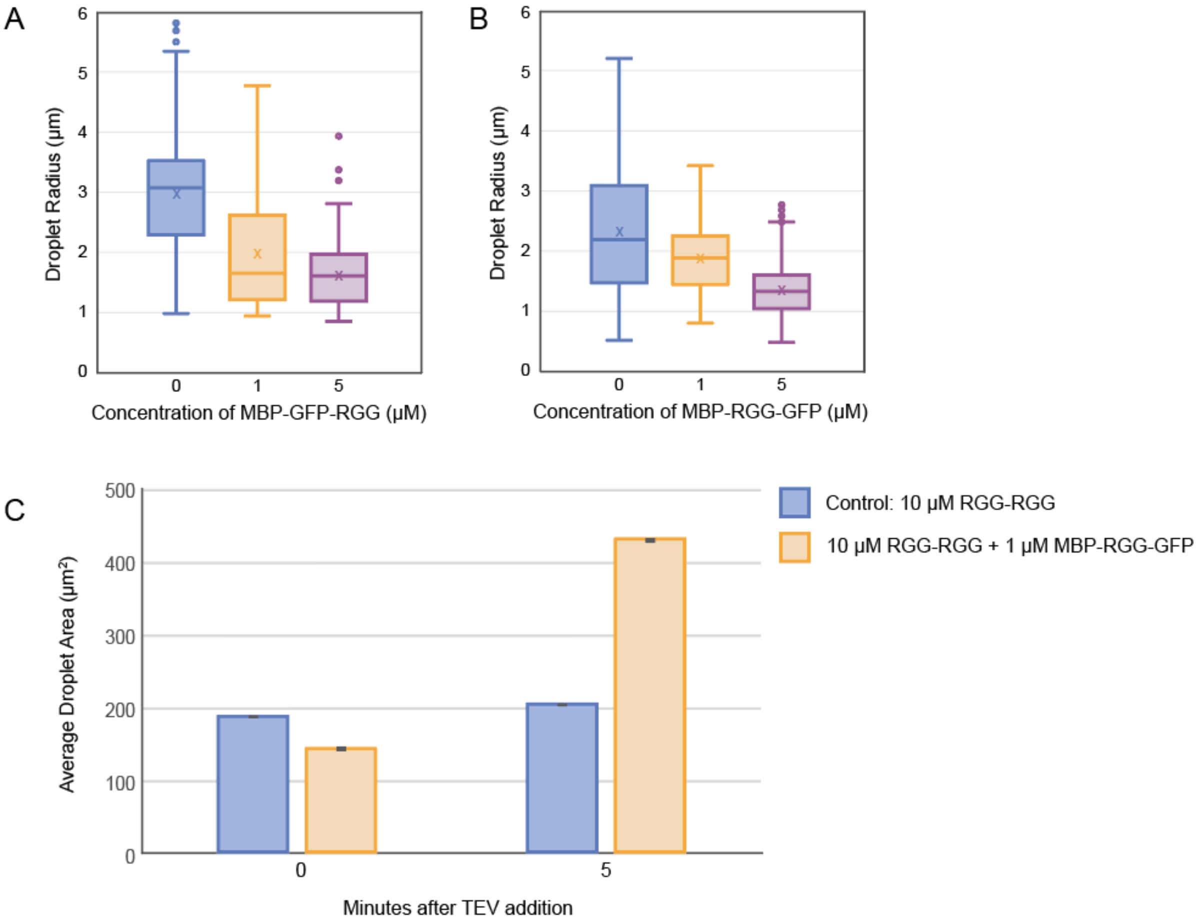


**Figure S4: Supporting data for control of RGG-RGG condensate size by surfactant proteins MBP-RGG-GFP and MBP-GFP-RGG. A)** Alternative representation of data shown in Fig. 3C using a box-and-whisker plot. Data shows increasing concentration of MBP-GFP-RGG results in condensates of smaller radii, on average. **B)** Alternative representation of data shown in Fig. 3F using a box-and-whisker plot. Data shows increasing concentration of MBP-RGG-GFP results in condensates of smaller radii, on average. **C)** Quantification of data shown in Fig. 3H. Comparison of average droplet size between RGG-RGG + 1 µM MBP-RGG-GFP (yellow) versus a control without a surfactant protein film (blue) upon adding TEV protease to both. TEV protease cleaves the MBP-RGG-GFP construct, releasing the surfactant protein film surrounding RGG-RGG droplets, resulting in rapid fusion events that increase droplet size. Error bars represent standard error of the mean and may be too small to see.

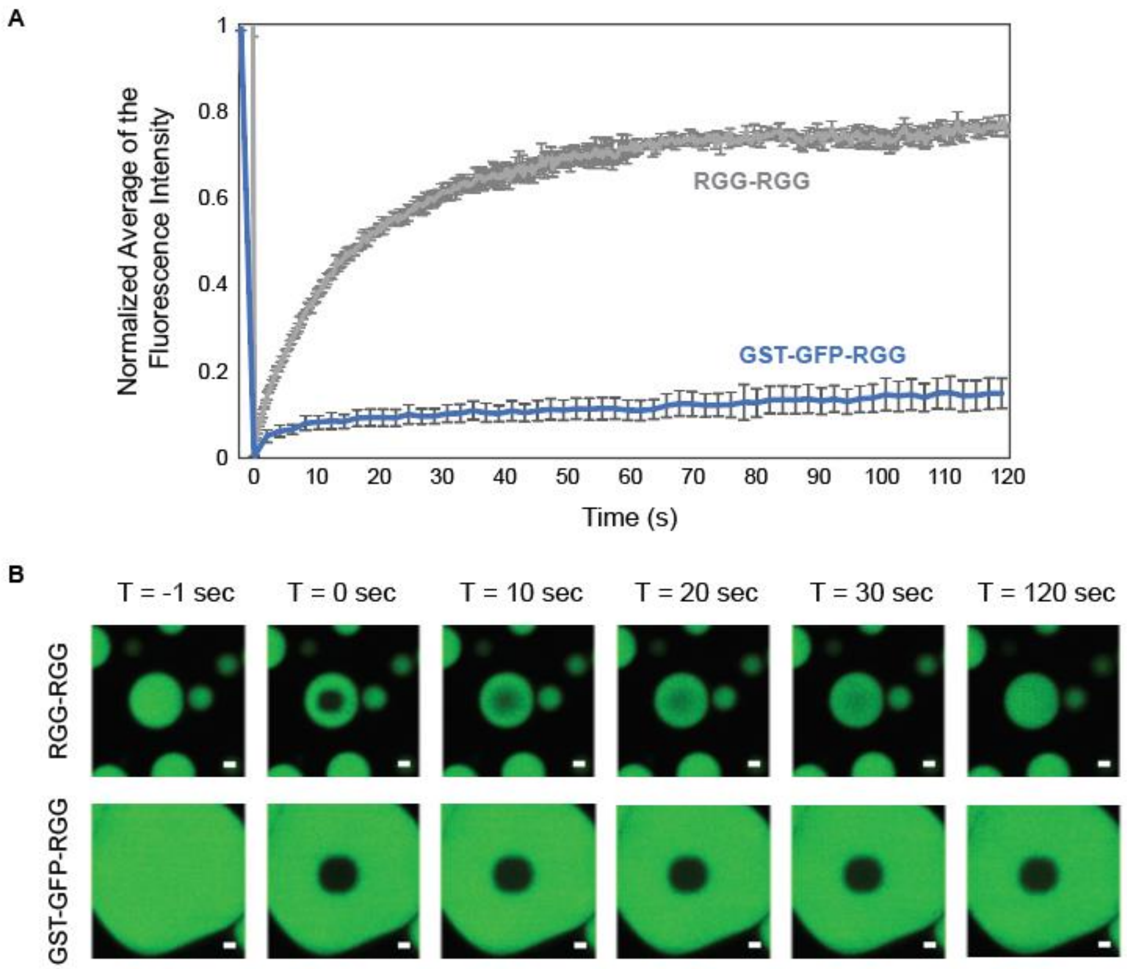


**Figure S5: Dynamics of GST-GFP-RGG vs. RGG-RGG condensates. A)** Normalized FRAP recovery curves of GST-GFP-RGG (blue) and RGG-RGG (grey). GST-GFP-RGG shows only a ~10% recovery of fluorescence after 2 minutes, indicating the phase is highly viscous or may be a gel-like material.  In comparison, RGG-RGG condensates recover quickly to around 80% of the original fluorescence within 1 minute; to allow for FRAP, RGG-RGG (9.5 μM) was mixed with RGG-GFP-RGG (0.5 μM). Error bars represent STD (n  = 3). **B)** Confocal fluorescence images from representative FRAP experiments show limited recovery of fluorescence within the bleached region of GST-GFP-RGG when compared to RGG-RGG over a period of 2 minutes. Scale bars, 1 μm.

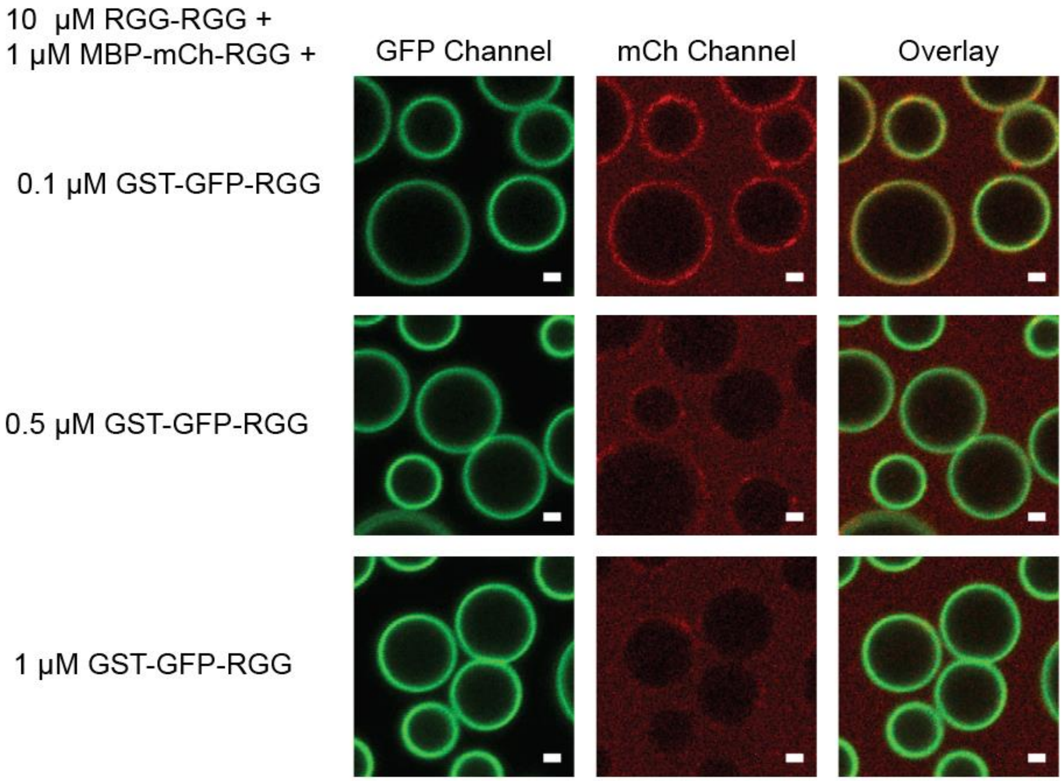


**Figure S6: Concentration-dependent competitive adsorption to the surface of RGG-RGG condensates when two amphiphilic proteins are introduced in different ratios.** Three samples, each with 10 μM RGG-RGG and 1 μM MBP-mCherry-RGG, were prepared with increasing concentrations of GST-GFP-RGG. Results show increasing competitive exclusion of MBP-mCherry-RGG from the condensate interface with increasing concentration of GST-GFP-RGG. Scale bars, 1 μm.

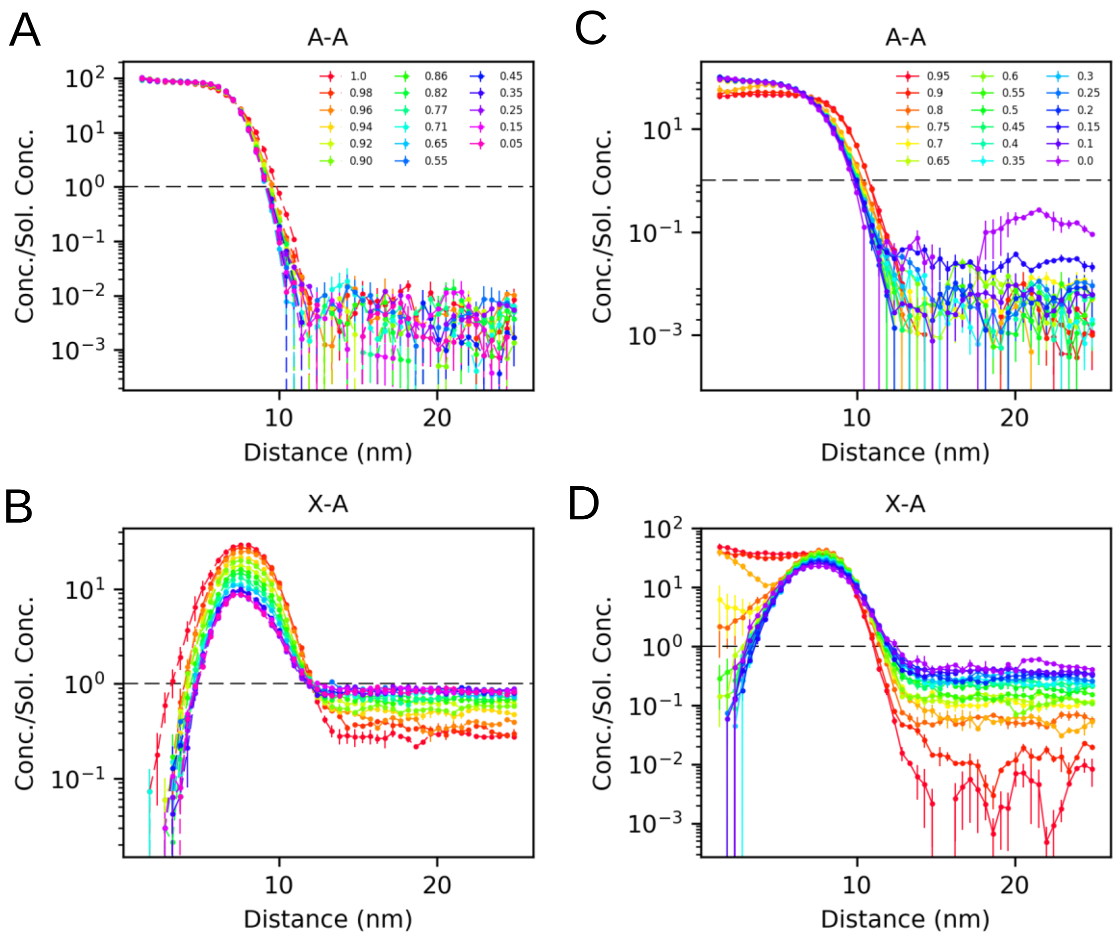


**Figure S7: Radial density profiles with respect to center of mass of A-A chains from coarse grained simulations of a binary (A-A/X-A) mixture for various heterotypic interaction strengths ($\epsilon_{AB}$) and homotypic interaction strengths ($\epsilon_{BB}$) show suitable interaction criteria for X-A.** A denotes the RGG domain and X denotes either the MBP or GST domains. Binary mixtures consisting of equal concentrations of both A-A and X-A chains were simulated at 298K for 30 nanoseconds with varying $\epsilon_{BB}$ with $\epsilon_{AB}$ fixed at 0.0 (A,B) and varying $\epsilon_{AB}$ with $\epsilon_{BB}$ fixed at 0.95 (C,D). Increasing $\epsilon_{BB}$ increases the density of X-A chains on the interface (B) whereas increasing $\epsilon_{AB}$ increases the concentration of X-A chains in the core of the condensate (D).

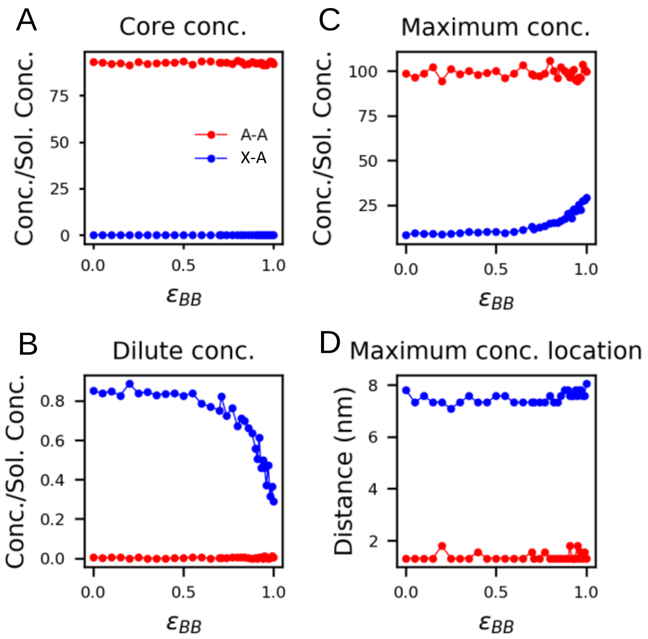


**Figure S8: Dilute phase concentration, core concentration, and maximum concentration and its location, from the density profiles for A-A and X-A chains in a binary mixture with varying $\epsilon_{BB}$ while $\epsilon_{AB}$ was fixed at 0.0 show the impact of homotypic X:X interactions on phase behavior.** A denotes the RGG domain and X denotes either the MBP or GST domains.

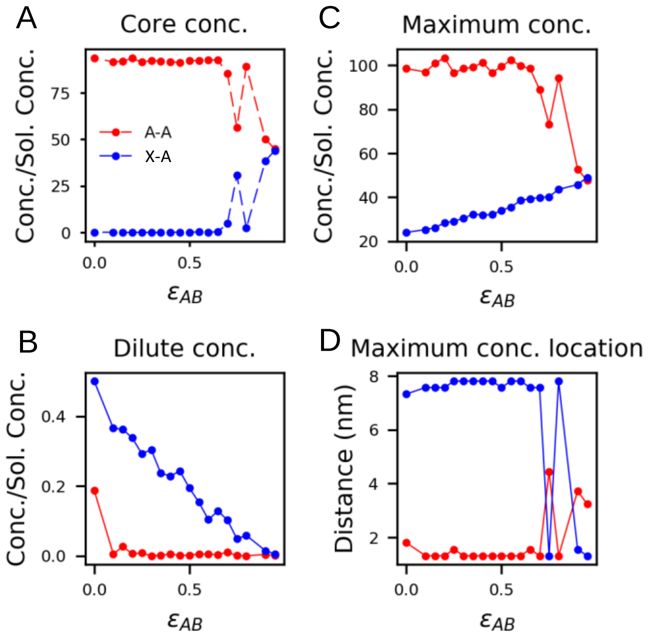


**Figure S9: Core concentration, dilute phase concentration, and maximum concentration and its location, from the density profile for A-A and X-A chains in a binary mixture with varying $\epsilon_{AB}$ while $\epsilon_{BB}$ was fixed at 0.95 show the impact of heterotypic X:A interactions on phase behavior.** A denotes the RGG domain and X denotes either the MBP or GST domains.

### 3. Bibliography

1. Schuster, B. S. *et al.* Identifying sequence perturbations to an intrinsically disordered protein that determine its phase-separation behavior. *Proc. Natl. Acad. Sci. U. S. A.* **117**, (2020).
